# Changes in body size with age do not follow the temperature-size rule

**DOI:** 10.1101/2025.09.02.673738

**Authors:** Jennifer S. Bigman, Lewis A.K. Barnett, James T. Thorson, Sean C. Anderson, Krista Oke, Kelly Kearney, Darren Pilcher, Wei Cheng, Esther Goldstein, Mary E. Matta, Kirstin K. Holsman, Lauren A. Rogers

## Abstract

The temperature size-rule is often described as a reduction in ectothermic body size with warming. This coincides with the expectation that faster growth under warming would lead to larger sizes early in life but smaller sizes later in life. Here, we use > 99,000 observations of weight-at-age of four commercially-important fishes over 25 years to ask whether changes in size and growth can be explained by temperature. We also examine whether oxygen, a key part of a proposed mechanism behind the temperature size-rule, explains patterns of weight-at-age better than temperature. Changes in weight-at-age over time were more related to temperature than oxygen but the effect of temperature on weight was small and did not vary by age, counter to the temperature size-rule. Importantly, these results were sensitive to how the relationship between weight-at-age and temperature was modeled. While the functional form (linear or polynomial) mattered little, how space was included in the model led to different conclusions regarding how temperature affects size and growth. Assuming that spatial and spatiotemporal random effects - those that account for the higher degree of similarity between observations collected closer in space and time - are shared across ages led to different results compared to allowing these effects to vary by age. Models with shared effects suggested weight for younger ages had positive relationships with temperature and negative relationships for older ages. However, these models provided spurious support for the temperature size-rule as they had less statistical support than models that allowed spatial effects to vary by age. Overall, our work highlights that relationships among size, growth, temperature, and oxygen may not be as straightforward as theory suggests and illustrates that modeling decisions can have a large effect on tests of ecological theory, and more broadly, our ability to understand biological responses to climate change.

## Introduction

One of the three universal responses of species to global change is shifts in body size (Daufresne et al., 2009; Gardner et al., 2011; Sheridan & Bickford, 2011). Such shifts are evident in both endothermic and ectothermic animals, plants, and bacteria, and species in both terrestrial and aquatic systems (Forster et al., 2012; Gardner et al., 2011; Martins et al., 2023). Both increases and decreases have been documented, but most organisms, and ultimately, populations, assemblages, and species, are getting smaller (Audzijonyte et al., 2020; Martins et al., 2023; Ohlberger, 2013). As body size is a central trait that influences almost all aspects of biology, reductions have countless implications for species and the systems in which they live (Andersen et al., 2016; Peters, 1986; Thorson et al., 2015). These implications span diverse biological and socioeconomic axes including ecosystem dynamics such as trophic relationships and other species interactions, as well as fisheries productivity (Audzijonyte et al., 2013; Baudron et al., 2014; Oke et al., 2020).

Increasing temperature is thought to be a major driver of declines in body size, particularly for ectotherms (Daufresne et al., 2009; Gardner et al., 2011; Sheridan & Bickford, 2011). The negative relationship between ectothermic body size and temperature is so widespread that it is formalized as the temperature-size rule. This pattern was originally described based on laboratory experiments in which individuals reared under warmer conditions grew faster to a smaller size-at-maturity but has since become much broader in scope (Atkinson, 1994; Audzijonyte et al., 2019, 2024). It is now commonly used to describe changes in size and growth with temperature in natural populations both within and across species, and is the subject of ecological theory concerned with global change (Audzijonyte et al., 2019, 2024; Verberk et al., 2021). For example, the temperature-size rule intersects life-history theory and body size optimization, metabolic theory, and theories of physiological performance including oxygen limitation (Brown et al., 2004; Essington et al., 2001; Marshall & White, 2019; Pörtner, 2010).

One proposed mechanism underlying the temperature-size rule that has received much attention is oxygen limitation, or the mismatch between oxygen supply and demand (Atkinson et al., 2006; Forster et al., 2012). This mismatch stems from the faster increase in oxygen demand (i.e., metabolic rate) compared to oxygen supply as temperature increases, which is thought to be exacerbated for larger organisms (Audzijonyte et al., 2019; Rubalcaba et al., 2020). Additionally, the magnitude of decline in body size is greater in aquatic versus terrestrial ectotherms, and coupled with the added challenges of extracting oxygen from water, supports the role of oxygen limitation behind the temperature size-rule (Forster et al., 2012; Rollinson & Rowe, 2018). More broadly, oxygen has been used in several mechanistic and phenomenological models to explain and predict widespread declines in maximum/asymptotic body size with warming (Bigman et al., 2023a; Cheung et al., 2013; Rubalcaba et al., 2020). However, the role of oxygen in the temperature-size rule remains debated and it is unclear whether oxygen availability can explain and predict changes in body size and growth across levels of biological organization (Atkinson et al., 2022; Audzijonyte et al., 2019; Bigman et al., 2023b).

Many studies examining the temperature-size rule in natural systems, at least for fishes, often focus on whether the mean, maximum, or asymptotic body sizes of populations or species are smaller with warming and/or reduced oxygen availability (e.g., Baudron et al., 2014; Cheung et al., 2013; Van Rijn et al., 2017). There is less research addressing how size and growth trade off along the ontogenetic trajectory to give rise to the pattern of faster growth to a smaller maturation size, and presumably, smaller maximum or asymptotic size. Size-at-age, or the size of individuals at a given age, is a metric that has been used to examine both growth and body size in tandem in natural populations, and has been found to vary with temperature such that younger age classes are larger under warming but older age classes are smaller, matching theoretical predictions (Ikpewe et al., 2021; Neuheimer & Grønkjær, 2012; Ohlberger et al., 2018; Oke et al., 2022). Understanding how body size and growth trade off over the lifetime of a fish is particularly critical for commercially important species as individuals are recruited to the fishery at body sizes much smaller than maximum/asymptotic size (and sometimes smaller than size-at-maturity). Thus, changes in fisheries productivity (e.g., population biomass, maximum sustainable yield, yield-per-recruit) will result not just from reductions in maximum/asymptotic size but also changes in size and growth along the ontogenetic trajectory (Baudron et al., 2014; Free et al., 2019; Oke et al., 2022).

Commercially important fish species are generally data-rich, but working with these and other geo-referenced ecological data comes with added complexities (Cressie & Wikle, 2011; Thorson & Kristensen, 2024). Such data are often collected via monitoring surveys, where observations are collected at a specific place and season, often repeatedly over time. Such a sampling scheme results in observations being more similar to each other if collected closer in space and time, and this autocorrelation must be accounted for when modeling ecological processes using these data (Cressie & Wikle, 2011; Indivero et al., 2023; Thorson & Kristensen, 2024). Another layer of complexity is added when assessing patterns across ages (or any other group structure), such as those predicted by the temperature-size rule. For example, the geographic distribution of individuals in different age classes varies spatially and temporally, particularly for fishes. Coupled with the sampling of fish in space over time, it is likely that the degree of autocorrelation of observations in space and time differs across ages. Recent efforts to make spatiotemporal modeling frameworks more accessible, including those that allow random effects such as the spatial and spatiotemporal fields to be independent by age (or other grouping structure), have resulted in a rapid increase in the use of spatiotemporal modeling (Anderson et al., 2022; Thorson, 2019; Thorson et al., 2024). However, the structure of these complex models, including how spatial and spatiotemporal fields are specified, can bias results or lead to spurious conclusions and thus careful consideration of model structure is important (Commander et al., 2022; Dormann et al., 2007).

Here, we aim to understand how body size across ages varies with temperature and oxygen, and whether these seemingly simple relationships stemming from theory can further our understanding of how fisheries productivity may respond to environmental change. We focus on the weight-at-age of four ecologically and commercially important groundfish species in the eastern Bering Sea, a region that is rapidly changing yet supports some of the most valuable fisheries in the world. Specifically, we ask two questions: (i) does size-at-age vary over time and (ii) is variability in size-at-age explained by temperature or oxygen as expected from the temperature-size rule, where size-at-age of younger age classes have a positive relationship with temperature or oxygen and older age classes have a negative relationship. To answer these questions, we fit spatiotemporal generalized linear mixed models, which allow us to account for the fact that observations are collected in space over time through random effects implemented as two-dimensional Gaussian Markov random fields. Spatial fields account for the autocorrelation of observations collected in space and spatiotemporal fields account for the autocorrelation of observations collected in space and time and are estimated for each time step (here, year; Cressie & Wikle, 2011; Thorson & Kristensen, 2024). We explored different parameterizations of these random fields to understand the implications of modeling decisions on our results. Specifically, we examined and compared models with (a) a spatial field that is shared across ages (‘shared spatial field’), (b) one spatial field and spatiotemporal fields for each year that are shared across ages (‘shared spatial and spatiotemporal fields”), and (c) fully independent spatial and spatiotemporal fields by age (i.e., a spatial field estimated for each age class and a spatiotemporal field estimated for each age class and year, “age-specific fields”). Understanding how body size and growth are affected by temperature and oxygen, and whether model structure affects this understanding, will inform both ecological theory and fisheries management, and more broadly, help us predict biological responses to global change.

## Materials and methods

### Survey data

We collated weight and age data for individuals of four groundfish species: arrowtooth flounder (*Atheresthes stomias*), Pacific cod (*Gadus macrocephalus*), walleye pollock (*Gadus chalcogrammus*), and yellowfin sole (*Limanda aspera*). These four species were chosen based on data availability and for comparison with other studies examining demographic trends (e.g., Holsman et al. 2020, Oke et al. 2020). Specimens were collected in fishery-independent bottom trawl surveys in the eastern and northern Bering Sea and aged by the National Oceanic and Atmospheric Administration’s Alaska Fisheries Science Center (AFSC) using standardized procedures (for more information, see Matta & Kimura, 2012). The bottom trawl survey has historically focused on the eastern Bering Sea with periodic extensions to the northern Bering Sea, which have become more common in recent years (Markowitz et al., 2024). The survey also evolved over time in terms of the species and number of specimens collected for ageing, resulting in a different collection of years sampled for each species (Table S1).

We followed AFSC’s protocol to remove outliers from the age and weight dataset using a Bonferroni outlier test applied to the length-weight data (Rohan & O’Leary, 2024).

### Environmental variables

For temperature and oxygen, we obtained weekly averaged, spatially resolved values of bottom temperature and bottom oxygen (defined as the mean temperature over the bottom [deepest] 5 meters of the water column at each horizontal [latitude-longitude] location) from the hindcast of the Bering10K Regional Ocean Modeling System (hereafter, ‘Bering10K’; Kearney et al., 2020). This reanalysis-forced simulation spans the period of 1970–present (through 2024 at the time of this paper), and its domain covers the Bering Sea and northern Gulf of Alaska using a 10-km horizontal resolution and 30 terrain-following depth levels. It has demonstrated high skill in reproducing key biophysical dynamics of the Bering Sea, including the seasonal extent of sea ice, formation of the cold pool, and general circulation and stratification patterns across the shelf (Hermann et al., 2016; Kearney et al., 2020; Pilcher et al., 2019). We chose to use modeled temperature and oxygen as we wanted a temporally-averaged and spatially-resolved time series, and *in situ* measurements are only available during certain times of the year (i.e., during the bottom trawl survey) for a sparse selection of years. For each week of each year, we matched the location of each survey station to the closest grid cell from the Bering 10K using nearest neighbor matching (as implemented in the RANN package, Arya et al., 2019). We then averaged the temperature and oxygen values for the year preceding the survey (July of one year to June of the next year).

### Does weight-at-age change over time?

To assess whether weight-at-age changed over time, we fit spatiotemporal generalized linear mixed-effects models, with individual weight as the response variable and year as the (factor) predictor variable. Fitting one model per species (with age class as a fixed effect) rarely converged due to the low sample size of some factor levels (i.e., few samples of a given age class in a given year) and so we fit separate models for each age class of each species and removed those species/age combinations resulting in failed model convergence. All models accounted for spatial random fields and independent spatiotemporal random fields using the stochastic partial differential equation (SPDE) approximation to Gaussian random fields with Gaussian Markov random fields (Lindgren et al., 2011). These models were fitted using the R package *sdmTMB*, which interfaces input matrices and a finite-element mesh from the R package *fmesher* with Template Model Builder (TMB) (Anderson et al., 2022; Kristensen et al., 2016; Lindgren, 2024). Weight was log10-transformed prior to analyses.

### Does temperature or oxygen explain changes in size-at-age?

To determine whether changes in weight-at-age over time can be explained by temperature or oxygen, as suggested by the temperature size-rule, we fit and compared three types of models. The first type of model did not include an environmental covariate (i.e., temperature or oxygen). The second type of model included a ‘global’ effect of temperature or oxygen, where the effect of temperature and oxygen on weight is the same for all ages. The third type of model included an age-specific effect of temperature or oxygen on weight (i.e., an interaction between age and temperature or oxygen), where the effect of temperature or oxygen on weight is allowed to vary by age class. Because physiological and ecological relationships are complex, we explored three functional forms that may describe the relationship between weight and temperature/oxygen - linear, second order polynomial, and third order polynomial. All models had a fixed effect of age class to estimate age-specific intercepts. Weight was log10-transformed and temperature and oxygen were standardized (centered by the mean and scaled by the standard deviation) prior to analyses.

To explore how specifications of spatial and spatiotemporal fields affect our understanding of changes in weight-at-age with temperature and oxygen, we fit all models described above with three different random effect structures. First, we fit multivariate models where spatial and spatiotemporal fields (by year) were estimated for each age class, and each age class had its own residual variance (models with ‘age-specific fields’). These models are essentially equivalent to fitting a separate model for each age class. Second, we fit models with a shared spatial field and shared residual variance across ages, but with no spatiotemporal field (models with a ‘shared spatial field’). Finally, we fit models where age classes shared a spatial field, spatiotemporal fields for each year, and residual variance (models with ‘shared spatial and spatiotemporal fields’). To do so, we used the R package *tinyVAST*, which offers the flexibility to specify multivariate spatiotemporal models, and thus, random effect structures and error distributions by grouping structure (e.g., age, size, species; Thorson et al., 2024; Thorson & Barnett, 2017). The shared spatial field models are similar to fitting a generalized additive model (GAM) or generalized linear model (GLM) with an interaction (e.g., tensor product) of latitude and longitude to account for space, an approach used to ask similar questions (e.g., Oke et al., 2022). The shared spatial and spatiotemporal fields models are most similar to fitting a model in *sdmTMB* with both spatial and spatiotemporal effects, as in this case, the spatial and spatiotemporal random fields would be shared across ages (Anderson et al., 2022).

We first fitted all models described above using restricted maximum likelihood (REML) estimation and then used (marginal) Akaike Information Criteria (AIC) to identify the most supported random effect structure. We then re-fit models with the most supported random effect structure with maximum likelihood (ML) estimation and used AIC to understand (1) whether temperature or oxygen better explained patterns of weight-at-age, (2) whether the relationship between temperature or oxygen and weight was age-specific (i.e., is the interaction important?), and (3) which functional form — linear, second order polynomial, or third order polynomial — best describes the relationship between weight and temperature or oxygen. Finally, to understand how the different random effect structures affected these three questions, we also re-fit models with the two suboptimal random effect structures with ML estimation (to facilitate comparison of predicted trends). In total, we fit 312 spatiotemporal models combined for the four fish species to test the effects of temperature and oxygen on weight-at-age (see Table S3, which reports all models run using one likelihood estimation [REML] for one species). Specifically, for each species, we fit:

i. models without an environmental covariate (“no covariate model”) with three different spatial and spatiotemporal random effect structures (shared spatial fields, shared spatial and spatiotemporal fields, and age-specific fields) both using ML (facilitates comparison of fixed effects) and REML (facilitates comparison of random effects) estimation (n = 6 models for each species, 24 models total),
ii. models with a global effect of *temperature* on weight as a linear, second order polynomial, and third order polynomial effect, each with three spatial and spatiotemporal random effect structures and with both ML and REML estimation (n = 18 models for each species, 72 models total),
iii. models with an age-specific effect of *temperature* on weight as a linear, second order polynomial, and third order polynomial effect, each with three spatial and spatiotemporal random effect structures and with both ML and REML estimation (n = 18 models for each species, 72 models total),
iv. models with a global effect of *oxygen* on weight as a linear, 2nd order polynomial, or 3rd order polynomial effect, each with three spatial and spatiotemporal random effect structures and with both ML and REML estimation (n = 18 models for each species, 72 models total), and
v. models with an age-specific effect of *oxygen* on weight as a linear, 2nd order polynomial, or 3rd order polynomial effect, each with three spatial and spatiotemporal random effect structures and with both ML and REML estimation (n = 18 models for each species, 72 models total).

A change in Pacific cod otolith age determination criteria has been documented wherein ages prior to 2008 were generally overestimated (Barbeaux et al., 2018). To ensure this did not affect our model results for Pacific cod specifically, we refit all models with environmental covariates (temperature and oxygen) to data from years 2008 and forward for Pacific cod. We then compared the top model with the full dataset to that of the dataset 2008 and forward.

All data and code are available at (https://github.com/jennybigman/size-at-age).

## Results

### Changes in weight-at-age over time

Weight-at-age was highly variable over time for all four species (Figure 1). While many ages had enough data per year to assess a change in weight over time, some did not. We could assess whether weight-at-age changed for ages 2 - 6 and age 8 for arrowtooth flounder, ages 1 - 7 of Pacific cod, ages 1 - 14 for walleye pollock, and ages 3 - 11 and age 28 for yellowfin sole.

**Figure 1.**
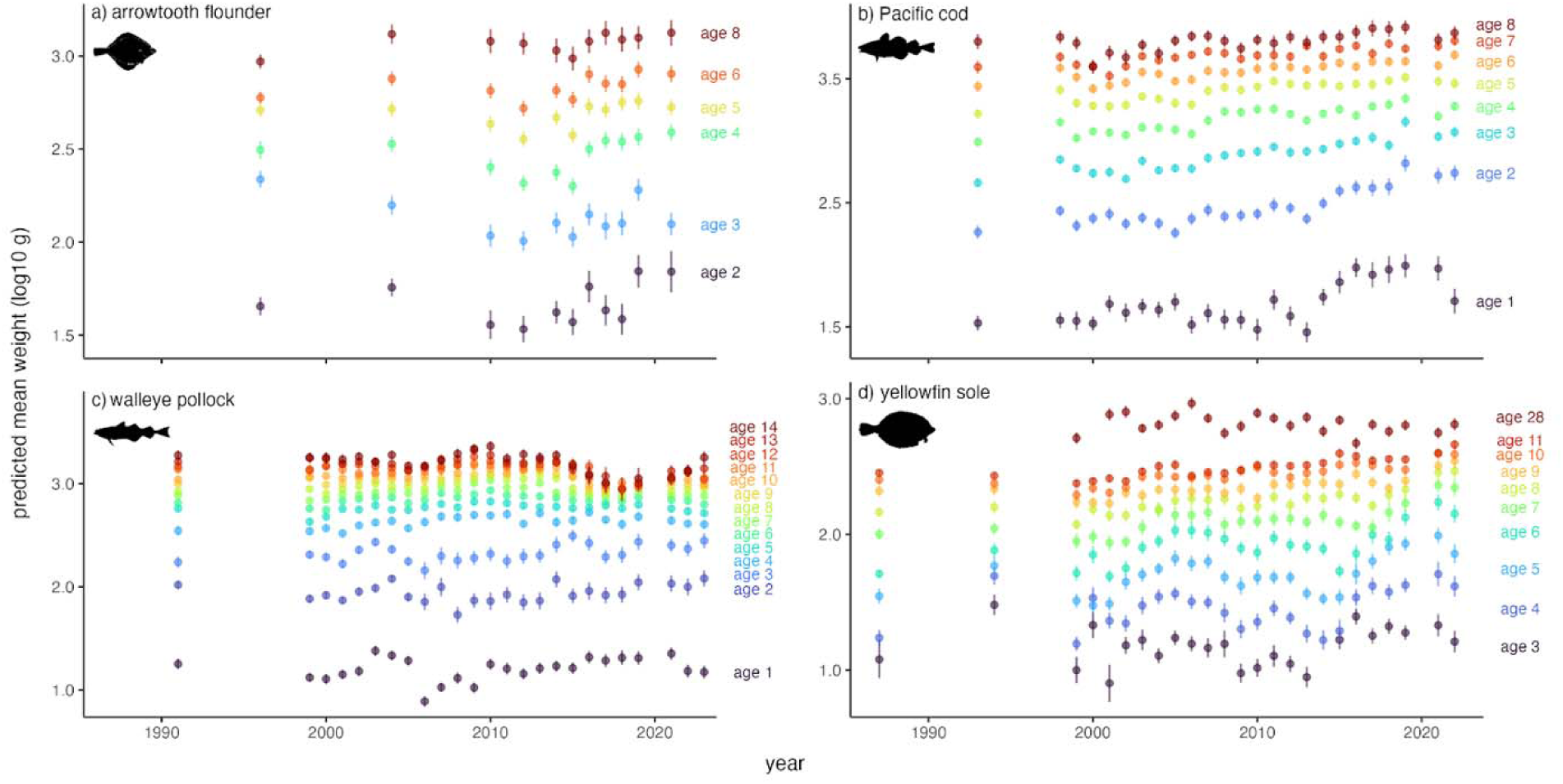
Weight-at-age was variable over time for all four species. Change in mean weight-at-age over time for Arrowtooth flounder (top left), Pacific cod (top right), Walleye pollock (bottom left), and Yellowfin sole (bottom right). Colored circles are the mean predicted weight-at-age and error bars are one standard error around the mean (see text). Color indicates age with labels insetted in each panel. Note the y-axis (the predicted mean weight [log10 g]) differs for each panel.

Although no significant positive or negative trend in weight-at-age over time existed for any species or age over the span of the survey period (Figure S1), weight for many ages of arrowtooth flounder, Pacific cod, and yellowfin sole appeared to be increasing in recent years. For Pacific cod, weight started increasing around the early to mid 2000s for most ages. Younger ages had a sharper increase around the mid 2010s (age 1 through age 4) and older ages showed a steadier increase since the early 2000s (Figure 1). The phase shift in weight-at-age for Pacific cod observed between 2007 and 2008 is likely at least partially a result of ageing bias (Barbeaux et al. 2018; although this ageing bias did not affect our model results, see below). Weight-at-age for arrowtooth flounder appears to be increasing since the mid 2010s, although this species has the shortest time series. Weight-at-age was more variable for walleye pollock and yellowfin sole. For walleye pollock, weight-at-age increased in the late 2000s for younger ages (age 1 - age 3) but for older ages, appears to have decreased since the late 2000s. However, weight-at-age for some older age classes seems to be stable since around 2016 (Figure 1). Weight-at-age for yellowfin sole increased and then subsequently decreased from 2000 to the late 2010s for ages 3 through 6; however, for older ages, weight-at-age steadily increased from the 2000s until present. Many ages had a sharp increase in weight-at-age starting around 2021, however, no data were collected in 2020.

### Does temperature or oxygen explain changes in weight-at-age over time?

To understand whether temperature or oxygen best explains patterns of size-at-age over time and whether these relationships follow our expectation from the temperature-size rule, we first identify the most supported random effect structure and then use those models to understand how temperature and oxygen affect patterns of size-at-age.

Based on AIC and residual patterns over time and space, the models with age-specific fields — those that allowed independent spatial and spatiotemporal random fields by age class (and age-specific residual variances) were strongly supported over models with a shared spatial field and models with a shared spatial field and shared spatiotemporal fields (Tables S2-S5, Figure 2). The AIC values of models with the same fixed effects but with different random effect structure were systematically lower for the models with age-specific fields. When mapping the residuals, there were clear patterns in space and time for models with shared fields (Figure 2, Figures S6-S9). For models with a shared spatial field, the residuals clustered in space and time such that certain areas and years had larger than expected weights and certain areas and years had smaller than expected weights. These patterns were also true for the models with shared spatial and spatiotemporal fields, although with some improvement. Yet when examining residuals spatially and temporally for the models with age-specific fields, much of the patterning in residuals in space and time that was evident in the other models was reduced (Figure 2, Figures S6-S9). Together, these results suggest that the models with age-specific fields — those with spatial and spatiotemporal fields that are allowed to vary by age class — are strongly supported over models that share the spatial or spatial and spatiotemporal fields across ages. Thus, we use the models with age-specific fields to understand how weight-at-age has changed over time with temperature and oxygen.

**Figure 2.**
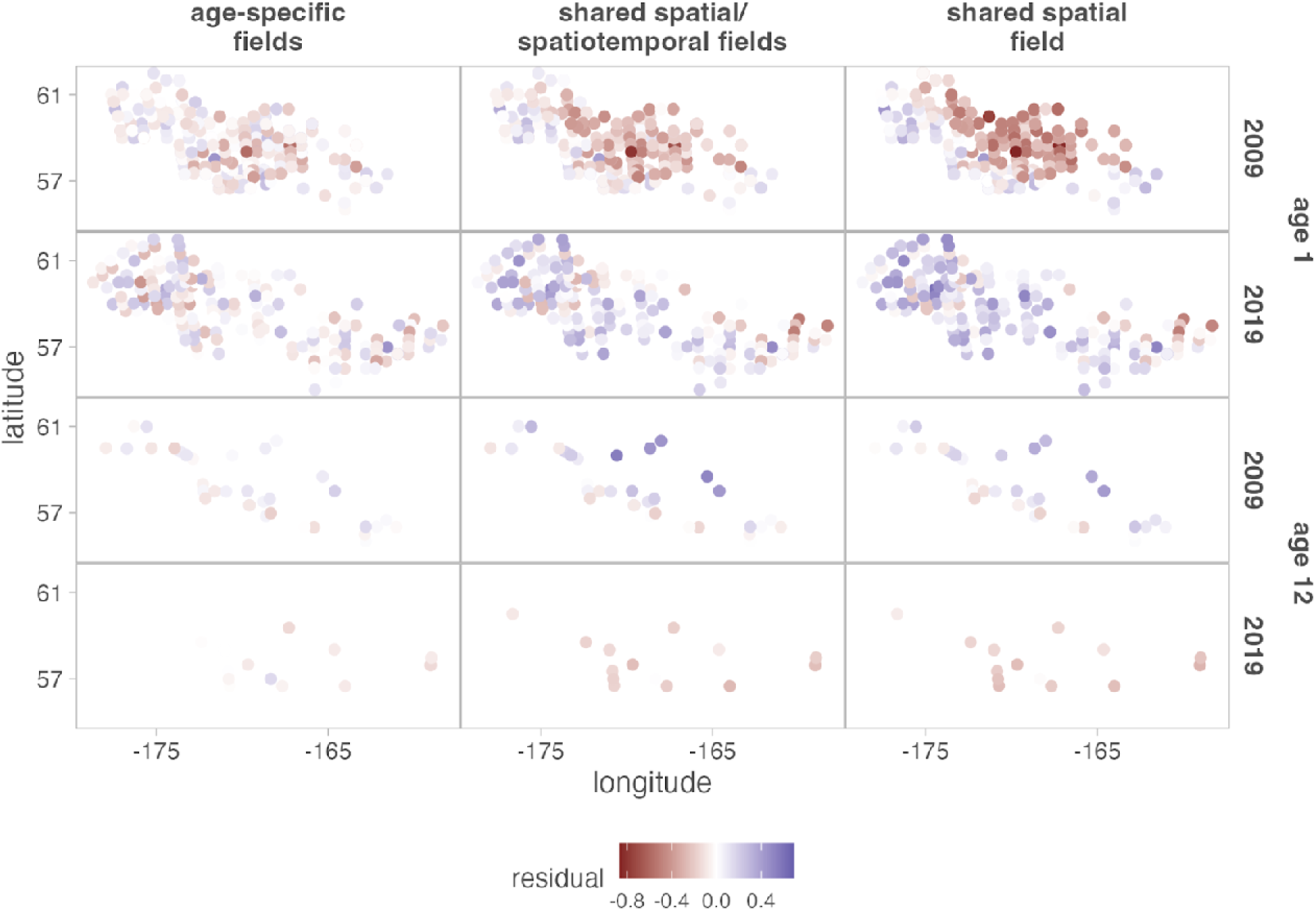
Models with independent spatial and spatiotemporal fields by age class have fewer residual patterns over space and time compared to models with shared effects. Comparison of residuals over space and time for two distant age classes (1 and 12, rows) for two distant years (2009, 2019, rows) among models with the same fixed effects and functional form but different specification of the random fields: independent spatial and spatiotemporal random fields across ages (first column), shared spatial and spatiotemporal effects across ages (middle column), and shared spatial effects across ages with no spatiotemporal effect (right column). Residuals shown here are for Walleye pollock and calculated from a model fitted to weight with a fixed effect of age class as a factor, an effect of temperature, and an interaction between age class and temperature, where the effect of temperature on weight is a second order polynomial (i.e., the most supported model structure for Walleye pollock; see Table S3). Figures S9-S12 show maps of residuals for all age classes for all years for all species.

Temperature was a better predictor of weight compared to oxygen for arrowtooth flounder and walleye pollock, as models with temperature had more support (Table S6, S8). The most supported models for Pacific cod (whose results did not differ between the top model fitted to the full dataset [all years] and a trimmed dataset [2008 and forward]; Figure S10) and yellowfin sole also included temperature, but out of all models within two AIC units, one model included oxygen (Table S7, S9). For Pacific cod, four models were within two AIC units from the most supported model and of the five total models, four included temperature and not oxygen. Likewise, three models were within two units of AIC of the top model for yellowfin sole and of the four total, three included temperature and not oxygen. Although models with temperature were more supported than models without, the effect size of temperature on weight was small and nonlinear for all species (Figures S2-S5).

Counter to the temperature-size rule, which suggests stronger positive correlations with temperature and size at younger ages, the relationship between weight and temperature was not age-specific, meaning that the most supported models did not have an interaction between age and temperature (Figure 3, Figures S2-S5; Tables S6-S9). This was true for many of the top-ranked models for all species (i.e., the top three to seven models depending on species did not include an interaction between temperature and age class, Tables S6-S9).

**Figure 3.**
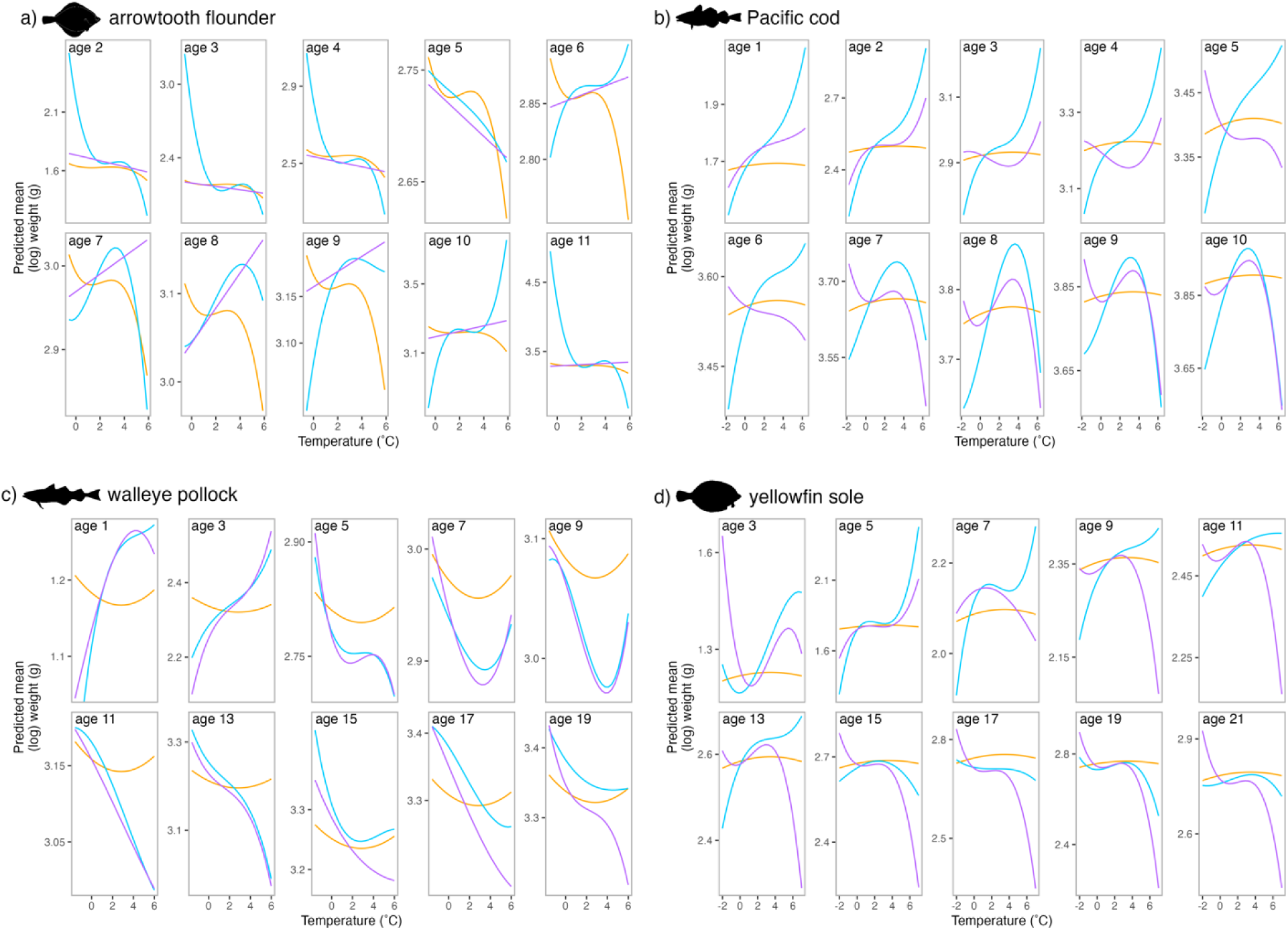
Model structure affected our understanding of how temperature affects weight-at-age: models with shared spatial and spatiotemporal fields provide spurious support for the temperature-size rule. Predictions of weight from temperature from three spatiotemporal models: (i) age-specific spatial and spatiotemporal fields (orange lines), (ii) shared spatial field and spatiotemporal fields across ages (purple lines), and (iii) shared spatial field across ages with no spatiotemporal fields (blue lines) for (a) arrowtooth flounder, (b) Pacific cod, (c) walleye pollock, and (d) yellowfin sole. Each panel corresponds to a separate age class (only 10 ages shown for each species; labeled in the upper left-hand side of the panel). Predictions for the three models with different random effect structures (orange, purple and blue lines) were generated from the most supported model for each random effect structure (i.e., most supported models for each random effect structure in Tables S2-S5). Note the y-axis (the predicted mean weight [log10 g]) differs for each panel.

There were two notable differences between the models with age-specific fields and those with shared fields (both the shared spatial field models and the shared spatial and spatiotemporal fields models) that are important to consider in the context of understanding whether temperature or oxygen can explain variation in size-at-age. First, the models with shared fields (the shared spatial field models and the shared spatial and spatiotemporal fields models) suggested that the relationship between weight and temperature and weight and oxygen differed by age class (Figure 3, Tables S10-S13). This was not the case for the most supported models overall—those with age-specific fields—as these models all found support for an additive effect of temperature, or one that is the same for all ages (Figure 3, Tables S10-S13). Second, the most supported models with shared spatial fields and the shared spatial and spatiotemporal fields suggested that the relationship between weight and temperature followed expectations from the temperature-size rule, where younger ages prior to maturity had generally positive relationships between weight and temperature and older ages after maturity had generally negative relationships (Figure 3). Again, this result was not found in the most supported models overall — those with age-specific fields, which did not find support for age-specific relationships between weight and temperature and weight and oxygen (Figure 3; Tables S6-S9).

Although the functional form of many of the most-supported models was either second or third order polynomial, comparison based on AIC indicated similar support for different functional forms (Figure 3, Tables S6-S9). This was especially true for Pacific cod and yellowfin sole, where the five and four highest-ranked models, respectively, were all within two AIC units of each other and described the relationship between weight and temperature as linear or as a second or third order polynomial (Tables S7, S9). For arrowtooth flounder and walleye pollock, the top two-ranked models had the lowest AIC, but were within two units of each other and described the weight-temperature relationship as a third order and second order polynomial (Tables S6, S8).

## Discussion

Our work suggests that while temperature explains changes in size and growth over time, this effect is small and not age-specific. Thus, we did not find support for the temperature-size rule — the pattern whereby warming is expected to lead to faster growth and larger sizes for younger age classes but smaller sizes for older age classes. Instead, we found more support for the same effect of temperature on size for all age classes. Although temperature did better at explaining historical patterns of weight-at-age compared to oxygen, the effects of both environmental covariates on weight were small, likely due to the small overall change in weight-at-age over time for all species. We also found that model structure strongly affected our understanding of the relationships between weight and temperature. Models that forced spatial and/or spatiotemporal random fields to be shared across ages supported the temperature size rule, where younger ages had positive relationships between weight and temperature and older ages had negative relationships. However, models with shared fields (either shared spatial or shared spatial and spatiotemporal fields) had less support compared to those with age-specific effects, which did not support predictions of the temperature-size rule as all age classes had the same temperature effect on weight for all four species. Collectively, our results indicate that the effect of temperature and oxygen on size and growth may not be as simple as suggested by ecological theory and highlight the complexities of modeling such relationships from fisheries data.

We found that although the effects of both environmental variables on weight were small, temperature generally had more support in explaining historical patterns of weight-at-age compared to oxygen, especially for two of the four species (walleye pollock and arrowtooth flounder). This result suggests that oxygen is unlikely to be a universal driver of changes in size and growth and contrasts with explanations that focus on oxygen’s central role behind the temperature-size rule (Atkinson et al., 2006; Forster et al., 2012; Pörtner, 2010). For example, the Gill Oxygen Limitation Theory (GOLT) argues that growth and size of water-breathing ectotherms is constrained by oxygen supply at the gills, which, as an organism increases in size, will be outpaced by the demand for oxygen for growth and other processes that require energy from aerobic metabolism (Pauly, 2021). Despite the GOLT and the wider body of literature suggesting oxygen may be behind the temperature-size rule, oxygen-related explanations have generally received mixed support when tested (Audzijonyte et al., 2019; Verberk et al., 2021), as found here. For example, Rubalcaba et al. (2020) found that the mismatch between oxygen supply and demand, termed oxygen limitation, limits aerobic scope (and presumably growth and size) in larger, more active fish in warmer waters. However, other work has identified that in fishes, oxygen supply does not strongly relate to maximum size and growth (Bigman et al., 2023a, 2023b; Wong et al., 2021). While we do not attempt to disentangle the many proposed mechanisms to explain the temperature-size rule (e.g., prey availability, life history theory, cell size, etc. [Angilletta et al., 2004; Audzijonyte et al., 2019; Verberk et al., 2021]) our work indicates that oxygen is unlikely to invariably explain changes in size and growth with temperature and other explanations are likely important. Whether the availability of oxygen affects life history and demographic processes may come down to local oxygen availability; for example, oxygen availability is likely more limiting in other systems.

The effect of temperature on weight was consistently small across ages and species. Both strong and weak effects of temperature on size and growth have been documented in natural populations (Martins et al., 2023; Solokas et al., 2023; Verberk et al., 2021). Audzijonyte et al. (2020) found increases, decreases, and no change in body size when examining trends in size across spatial (latitudinal) and temporal changes in temperature in 335 unexploited coastal reef fishes. Although over half of the species were smaller in warmer waters, a high percentage (45%) were larger. This pattern seemed to be related to the maximum size of a species: smaller-bodied species (those with smaller maximum sizes) were more likely to become smaller in warmer waters (Audzijonyte et al., 2020). Other studies have identified opposite patterns between maximum size and the strength of changes in size with temperature. For instance, Horne et al., (2015) found that temperature-size responses were more negative in larger-bodied arthropods and Huss et al., (2019) found that only smaller-bodied species had increased growth under warmer temperatures. The strength and direction of temperature-size relationships in our study do not seem to relate to maximum size as the two largest species had opposite responses of size and temperature, but a general pattern is hard to discern across four species. In part, these different responses to temperature may be related to responses of growth to temperature as growth increases with increasing temperature up to a point (e.g., optimum) and decreases thereafter (Kingsolver & Woods, 2016; Ohlberger, 2013). If individuals are experiencing temperatures that are lower than their optimum temperature for growth, they may grow faster with warming (leading to larger sizes) and then growth may slow or stop when temperatures exceed those that are optimal. However, thermal sensitivities of growth are often measured on individuals in the laboratory under highly controlled thermal environments and such sensitivities may not reflect patterns at the population-or species-level scale or in natural environments as we observed here (Audzijonyte et al., 2024; Schulte et al., 2011; Van Denderen et al., 2020).

In contrast to the temperature-size rule, we did not find support for age-specific relationships between size and temperature (or oxygen), indicating that our models could not rule out that the effect of temperature and oxygen on weight was the same for all age classes. Despite mounting evidence for temperature-size responses in controlled laboratory settings, responses in natural populations remain mixed (Audzijonyte et al., 2020; Forster et al., 2012; Verberk et al., 2021). In the laboratory, size and growth are often measured on an individual level and with relatively little error (Audzijonyte et al., 2024). Studies on natural populations often examine population-or species-level metrics of size and growth, metrics that are both observed (e.g., maximum size) and modeled (e.g., von Bertalanffy growth coefficient and asymptotic size), many of which naturally differ from laboratory-measured size and growth (Audzijonyte et al., 2024; Verberk et al., 2021). Additionally, size and growth data from natural populations come from individuals that have survived size-dependent fishing and natural mortality, forming a biased sample. When assessing the effect of temperature on size and growth, it is therefore important to consider what the expected relationship with temperature (or other predictors) may be in a particular setting or at a particular biological level (individual, population, species). Here, we expanded explorations of the temperature-size rule by looking beyond changes in the mean/asymptotic/maximum size of a population or species by using weight-at-age and testing whether patterns across ages matched theoretical expectations. However, size-at-age (both weight-and length-at-age) is still an imperfect metric as other processes aside from growth contribute to the realized size of a given age class of a population. Identifying metrics of both size and growth that facilitate detection of changes with temperature in natural populations remains a challenge.

We found that our understanding of how size and growth vary with temperature (or oxygen) across ages was sensitive to model structure, specifically how space and time were accounted for. When we compared models with shared versus age-specific spatial and spatiotemporal random effects, we saw contrasting results as models with shared effects supported temperature-size rule predictions. Similarly, a previous study examining changes in weight-at-age with temperature for walleye pollock in the Bering Sea — the same population included in this study — found support for the temperature-size rule as the relationship between weight and temperature was positive early in life and negative later in life (Oke et al., 2022). While that study used different temperature data than used here (regional mean observed sea surface temperature versus our spatially-explicit bottom temperature from a well-validated ocean model), the model formulation was most similar to our model with a shared spatial effect across all age classes (and no spatiotemporal effect), which yielded very similar results. However, we found that models with age-specific spatial and spatiotemporal fields were a better fit to the data. This is unsurprising as ecology and physiology (e.g., distribution, thermal limits) are known to change across ontogeny. The task of modeling these dynamics over time and space, and more broadly, the effects of different spatial and spatiotemporal effects specifications, is an area of active research (Hodges & Reich, 2010; Thorson et al., 2023; Ward et al., 2022). As found here, Commander et al. (2022) highlighted that how spatial and spatiotemporal effects are specified can change model inference. Predictions of biomass and the center of gravity (i.e., the mean of the distribution) for sablefish off the West Coast of the United States differed depending on whether spatial or spatiotemporal random effects were included, as well as whether spatiotemporal effects were independent or correlated through time (Commander et al., 2022). However, others argue that spatial and spatiotemporal random effects, as implemented in spatiotemporal modeling frameworks such as *sdmTMB* and *tinyVAST*, may be too flexible and soak up a lot of variation, requiring a lot of data to detect small effects (e.g., Hodges & Reich, 2010). Here, we worked with some of the most data-rich datasets on size-at-age globally and still detected small effects after accounting for spatial and spatiotemporal effects. Future work could extend our exploration of how random effects influence fixed effects through methods such as simulation. Given the importance of spatiotemporal models in marine resource management coupled with the explosion of their use as software becomes more user-friendly, it is crucial to fully explore the effects of model structure and how spatial and spatiotemporal effects alter results before drawing conclusions and using model output in fisheries management.

Although our work advances the exploration of shifts in size and growth with temperature in natural populations, many challenges in understanding and predicting this response remain. First, disentangling the dynamics among size, growth, and temperature is difficult for numerous reasons, many of which are interrelated. A general consensus regarding the temperature-size rule or a consistent pattern across species, systems, and settings is lacking (Audzijonyte et al., 2019, 2024; Verberk et al., 2021). Whether this is due to the use of different metrics for growth and size, examining changes in size and growth in different life stages, scaling up responses at the individual-level to the population-or species-level, understanding how responses in the laboratory translate to those in natural populations, modeling approaches, or other reasons, it makes it difficult to identify the underlying mechanism by which temperature affects size and growth. In turn, the lack of a mechanistic understanding hinders our ability to predict how warming will affect size and growth of fishes. Second, it is important to consider how temperature (or any other predictor) is summarized temporally and spatially for modeling. While size-temperature responses are evident at both coarse and fine spatial and temporal scales (suggesting that these relationships may be robust to the different ways temperature is spatially and temporally summarized; Cheung et al., 2013; Lindmark et al., 2024), future work could identify best practices for summarizing time series of environmental variables that vary over space and time in relation to the biological response being studied. This would be especially valuable for species that have ontogenetic shifts in depth or habitat use and for working with data from fisheries surveys, as catchability may differ by age class. Fourth, it may be that other abiotic or biotic factors more strongly affect size and growth compared to temperature. For example, competition, food availability, and demographic considerations (e.g., density dependence) are known to affect size and growth in both natural populations and experimental settings, and for harvested populations, fishing often results in similar changes in size and growth (Angilletta, Jr., & Dunham, 2003; Ohlberger et al., 2023; Waples & Audzijonyte, 2016). It is likely that interacting ecological and physiological processes, not just temperature, give rise to changes in size across levels of biological organization (Audzijonyte et al., 2019; Gårdmark & Huss, 2020). Last, statistical modeling approaches to detect changes in size or growth with temperature in natural populations must be suited to the available data, which are often inherently complex. As shown here, spatiotemporal modeling frameworks, which help us account for the spatial and spatiotemporal autocorrelation in geo-referenced data, can be specified in multiple ways, and seemingly small choices can have a large effect on inference. A shared understanding of the challenges and considerations in working across systems and scales, both regions and in the laboratory versus natural populations (e.g., how should a specific metric of size or growth change with temperature in a specific region or functional group), would go far in advancing the field. A first step in this direction would be to outline best practices for designing experiments and statistical models that aim to detect the effects of temperature on size and growth.

Commercially important fish species pose both opportunities and obstacles to studying responses to climate change (e.g., Hobday and Evans 2013). Such species are generally data-rich compared to unharvested taxa and, thus, changes in size across the ontogenetic growth trajectory with temperature can be examined as done here. Yet, identifying and understanding responses to environmental change in harvested taxa remains challenging. In addition to the complexity of modeling these types of data, accounting for the effect of fishing, which is known to truncate age- and size-structure, is a central challenge when working with data from exploited populations (Audzijonyte et al., 2016; Morrongiello et al., 2019; Waples & Audzijonyte, 2016). Another central challenge is not just understanding how species will respond to environmental change, but predicting how they will change. Our work is a step in this direction as we identify that while temperature (and oxygen) does relate to changes in size, this relationship is not consistent across species nor likely strong enough to be useful for generating predictions of size-at-age from temperature. Future work in this area could include fishing effects and other processes that were not explored here (e.g. cohort and sex effects, Cheng et al., 2023; Stawitz et al., 2015), with the goal of identifying how best to predict size-at-age rather than testing an explicit theoretical prediction, as done here. As fisheries globally face an uncertain future due to continued and projected changing environmental conditions, it is critical to ensure the stability of future fisheries, as well as the livelihoods and economies they support. Doing so will require continued exploration into how climate change is affecting commercially important fish populations, ideally with the goal of predicting how distributions, biomass, and abundance will change in the future.

## Acknowledgments

We would like to thank the survey team (Groundfish Assessment Program) and the age and growth team (Age and Growth Program) at NOAA AFSC. We thank Christine Stawitz for reviewing this manuscript, and others at NOAA AFSC, NOAA Northwest Fisheries Science Center, and NOAA Office of Science and Technology (OST) for input and reviews. Finally, we thank the ACLIM group at NOAA AFSC for their comments and suggestions, and those who developed and implemented the Bering 10K model. JSB also acknowledges funding from the National Science Foundation, NSF grant number 2109411.

## Author contributions

JSB: conceptualization, data curation, formal analysis, funding acquisition, methodology, project administration, software, visualization, writing – original draft, writing – review and editing; LAKB: data curation, investigation, methodology, writing – review & editing; JTT: methodology, writing – review & editing; SCA: methodology, writing – review & editing; KO: writing – review & editing; KK: data curation, investigation, writing – review & editing; DP: data curation, investigation, writing – review & editing; WC: data curation, investigation, writing – review & editing; EG: investigation, writing – review & editing; MEM: investigation, writing – review & editing, KKH: data curation, writing – review & editing; LAR: methodology, supervision, writing – review & editing.

## References

Andersen, K. H., Berge, T., Gonçalves, R. J., Hartvig, M., Heuschele, J., Hylander, S., Jacobsen, N. S., Lindemann, C., Martens, E. A., Neuheimer, A. B., Olsson, K., Palacz, A., Prowe, A. E. F., Sainmont, J., Traving, S. J., Visser, A. W., Wadhwa, N., & Kiørboe, T. (2016). Characteristic Sizes of Life in the Oceans, from Bacteria to Whales. Annual Review of Marine Science, 8(1), 217–241. 10.1146/annurev-marine-122414-034144

Anderson, S. C., Ward, E. J., English, P. A., Barnett, L. A. K., & Thorson, J. T. (2022). sdmTMB: An R Package for Fast, Flexible, and User-Friendly Generalized Linear Mixed Effects Models with Spatial and Spatiotemporal Random Fields. Ecology. 10.1101/2022.03.24.485545

Angilletta, Jr., M. J., & Dunham, A. E. (2003). The TemperaturelSize Rule in Ectotherms: Simple Evolutionary Explanations May Not Be General. The American Naturalist, 162(3), 332–342. 10.1086/377187

Angilletta, M. J., Steury, T. D., & Sears, M. (2004). Temperature, Growth Rate, and Body Size in Ectotherms: Fitting Pieces of a Life-History Puzzle. Integrative and Comparative Biology, 44(6), 498–509. 10.1093/icb/44.6.498

Arya, S., Kemp, S. E., & Jeffries, G. (2019). RANN: Fast Nearest Neighbour Search (Wraps ANN Library) using L2 metric (Version 2.6.1) [Computer software].

Atkinson, D. (1994). Temperature and organism size—A biological law for ectotherms?

Atkinson, D., Leighton, G., & Berenbrink, M. (2022). Controversial Roles of Oxygen in Organismal Responses to Climate Warming. The Biological Bulletin, 243(2), 207–219. 10.1086/722471

Atkinson, D., Morley, S. A., & Hughes, R. N. (2006). From cells to colonies: At what levels of body organization does the ‘temperaturelsize rule’ apply? Evolution & Development, 8(2), 202–214. 10.1111/j.1525-142X.2006.00090.x

Audzijonyte, A., Andersen, K. H., Atkinson, D., Bigman, J., Blanchard, J. L., Coghlan, A. R., Heather, F., Lindmark, M., Morrongiello, J. R., & Pauly, D. (2024). Temperature affects fish body sizes. Which sizes? Preprints. 10.22541/au.171813253.38400021/v1

Audzijonyte, A., Barneche, D. R., Baudron, A. R., Belmaker, J., Clark, T. D., Marshall, C. T., Morrongiello, J. R., & van Rijn, I. (2019). Is oxygen limitation in warming waters a valid mechanism to explain decreased body sizes in aquatic ectotherms? Global Ecology and Biogeography, 28(2), 64–77. 10.1111/geb.12847

Audzijonyte, A., Fulton, E., Haddon, M., Helidoniotis, F., Hobday, A. J., Kuparinen, A., Morrongiello, J., Smith, A. D., Upston, J., & Waples, R. S. (2016). Trends and management implications of human-influenced life-history changes in marine ectotherms. Fish and Fisheries, 17(4), 1005–1028. 10.1111/faf.12156

Audzijonyte, A., Kuparinen, A., Gorton, R., & Fulton, E. A. (2013). Ecological consequences of body size decline in harvested fish species: Positive feedback loops in trophic interactions amplify human impact. Biology Letters, 9(2), 20121103. 10.1098/rsbl.2012.1103

Audzijonyte, A., Richards, S. A., Stuart-Smith, R. D., Pecl, G., Edgar, G. J., Barrett, N. S., Payne, N., & Blanchard, J. L. (2020). Fish body sizes change with temperature but not all species shrink with warming. Nature Ecology & Evolution, 4(6), 809–814. 10.1038/s41559-020-1171-0

Barbeaux S et al. (2018) Assessment of the Pacific cod stock in the Gulf of Alaska. NPFMC, Anchorage, AK.

Baudron, A. R., Needle, C. L., Rijnsdorp, A. D., & Tara Marshall, C. (2014). Warming temperatures and smaller body sizes: Synchronous changes in growth of North Sea fishes. Global Change Biology, 20(4), 1023–1031. 10.1111/gcb.12514

Bigman, J. S., Wegner, N. C., & Dulvy, N. K. (2023a). Gills, growth and activity across fishes. Fish and Fisheries, 24(5), 730–743. 10.1111/faf.12757

Bigman, J. S., Wegner, N. C., & Dulvy, N. K. (2023b). Revisiting a central prediction of the Gill Oxygen Limitation Theory: Gill area index and growth performance. Fish and Fisheries, 24(3), 354–366. 10.1111/faf.12730

Brown, J. H., Gillooly, J. F., Allen, A. P., Savage, V. M., & West, G. B. (2004). Toward a metabolic theory of ecology. Ecology, 85(7), 1771–1789. 10.1890/03-9000

Cheng, M. Lh., Thorson, J. T., Ianelli, J. N., & Cunningham, C. J. (2023). Unlocking the triad of age, year, and cohort effects for stock assessment: Demonstration of a computationally efficient and reproducible framework using weight-at-age. Fisheries Research, 266, 106755. 10.1016/j.fishres.2023.106755

Cheung, W. W. L., Watson, R., & Pauly, D. (2013). Signature of ocean warming in global fisheries catch. Nature, 497(7449), 365–368. 10.1038/nature12156

Commander, C. J. C., Barnett, L. A. K., Ward, E. J., Anderson, S. C., & Essington, T. E. (2022). The shadow model: How and why small choices in spatially explicit species distribution models affect predictions. PeerJ, 10, e12783. 10.7717/peerj.12783

Cressie, N., & Wikle, C. K. (2011). Statistics for Spatio-Temporal Data. John Wiley & Sons.

Daufresne, M., Lengfellner, K., & Sommer, U. (2009). Global warming benefits the small in aquatic ecosystems. Proceedings of the National Academy of Sciences, 106(31), 12788–12793. 10.1073/pnas.0902080106

Dormann, C., M. McPherson, J., B. Araújo, M., Bivand, R., Bolliger, J., Carl, G., G. Davies, R., Hirzel, A., Jetz, W., Daniel Kissling, W., Kühn, I., Ohlemüller, R., R. Peres-Neto, P., Reineking, B., Schröder, B., M. Schurr, F., & Wilson, R. (2007). Methods to account for spatial autocorrelation in the analysis of species distributional data: A review. Ecography, 30(5), 609–628. 10.1111/j.2007.0906-7590.05171.x

Essington, T. E., Kitchell, J. F., & Walters, C. J. (2001). The von Bertalanffy growth function, bioenergetics, and the consumption rates of fish. Canadian Journal of Fisheries and Aquatic Sciences, 58(11), 2129–2138. 10.1139/f01-151

Forster, J., Hirst, A. G., & Atkinson, D. (2012). Warming-induced reductions in body size are greater in aquatic than terrestrial species. Proceedings of the National Academy of Sciences, 109(47), 19310–19314. 10.1073/pnas.1210460109

Free, C. M., Thorson, J. T., Pinsky, M. L., Oken, K. L., Wiedenmann, J., & Jensen, O. P. (2019). Impacts of historical warming on marine fisheries production. Science, 363(6430), 979– 983. 10.1126/science.aau1758

Gårdmark, A., & Huss, M. (2020). Individual variation and interactions explain food web responses to global warming. Philosophical Transactions of the Royal Society B: Biological Sciences, 375(1814), 20190449. 10.1098/rstb.2019.0449

Gardner, J. L., Peters, A., Kearney, M. R., Joseph, L., & Heinsohn, R. (2011). Declining body size: A third universal response to warming? Trends in Ecology & Evolution, 26(6), 285–291. 10.1016/j.tree.2011.03.005

Hermann, A. J., Gibson, G. A., Bond, N. A., Curchitser, E. N., Hedstrom, K., Cheng, W., Wang, M., Cokelet, E. D., Stabeno, P. J., & Aydin, K. (2016). Projected future biophysical states of the Bering Sea. Deep Sea Research Part II: Topical Studies in Oceanography, 134, 30–47. 10.1016/j.dsr2.2015.11.001

Hodges, J. S., & Reich, B. J. (2010). Adding Spatially-Correlated Errors Can Mess Up the Fixed Effect You Love. The American Statistician, 64(4), 325–334. 10.1198/tast.2010.10052

Horne, C. R., Hirst, Andrew. G., & Atkinson, D. (2015). Temperature-size responses match latitudinal-size clines in arthropods, revealing critical differences between aquatic and terrestrial species. Ecology Letters, 18(4), 327–335. 10.1111/ele.12413

Huss, M., Lindmark, M., Jacobson, P., Van Dorst, R. M., & Gårdmark, A. (2019). Experimental evidence of gradual size-dependent shifts in body size and growth of fish in response to warming. Global Change Biology, 25(7), 2285–2295. 10.1111/gcb.14637

Ikpewe, I. E., Baudron, A. R., Ponchon, A., & Fernandes, P. G. (2021). Bigger juveniles and smaller adults: Changes in fish size correlate with warming seas. Journal of Applied Ecology, 58(4), 847–856. 10.1111/1365-2664.13807

Indivero, J., Essington, T. E., Ianelli, J. N., & Thorson, J. T. (2023). Incorporating distribution shifts and spatio-temporal variation when estimating weight-at-age for stock assessments: A case study involving the Bering Sea pollock ( *Gadus chalcogrammus* ). ICES Journal of Marine Science, 80(2), 258–271. 10.1093/icesjms/fsac236

Kearney, K., Hermann, A., Cheng, W., Ortiz, I., & Aydin, K. (2020). A coupled pelagic–benthic– sympagic biogeochemical model for the Bering Sea: Documentation and validation of the BESTNPZ model (v2019.08.23) within a high-resolution regional ocean model. Geoscientific Model Development, 13(2), 597–650. 10.5194/gmd-13-597-2020

Kingsolver, J. G., & Woods, H. A. (2016). Beyond Thermal Performance Curves: Modeling Time-Dependent Effects of Thermal Stress on Ectotherm Growth Rates. The American Naturalist, 187(3), 283–294. 10.1086/684786

Kristensen, K., Nielsen, A., Berg, C. W., Skaug, H., & Bell, B. M. (2016). TMB: Automatic Differentiation and Laplace Approximation. Journal of Statistical Software, 70(5). 10.18637/jss.v070.i05

Lindgren, F. (2024). fmesher: Triangle meshes and related geometry tools (Version 0.2.0) [R].

Lindgren, F., Rue, H., & Lindström, J. (2011). An Explicit Link between Gaussian Fields and Gaussian Markov Random Fields: The Stochastic Partial Differential Equation Approach. Journal of the Royal Statistical Society Series B: Statistical Methodology, 73(4), 423–498. 10.1111/j.1467-9868.2011.00777.x

Lindmark, M., Ohlberger, J., & Gardmark, A. (2024). Non-linear growth-temperature relationship leads to opposite responses to warming in cold versus warm populations.

Markowitz, E. H., Dawson, E. J., Wasserman, S. N., Anderson, C. B., Rohan, S. K., Charriere, N. E., & Stevenson, D. E. (2024). Results of the 2023 eastern and northern Bering Sea continental shelf bottom trawl survey of groundfish and invertebrate fauna (NOAA Technical Memorandum No. NMFS-AFSC-487).

Marshall, D. J., & White, C. R. (2019). Aquatic Life History Trajectories Are Shaped by Selection, Not Oxygen Limitation. Trends in Ecology & Evolution, 34(3), 182–184. 10.1016/j.tree.2018.12.015

Martins, I. S., Schrodt, F., Blowes, S. A., Bates, A. E., Bjorkman, A. D., Brambilla, V., Carvajal-Quintero, J., Chow, C. F. Y., Daskalova, G. N., Edwards, K., Eisenhauer, N., Field, R., Fontrodona-Eslava, A., Henn, J. J., Van Klink, R., Madin, J. S., Magurran, A. E., McWilliam, M., Moyes, F., … Dornelas, M. (2023). Widespread shifts in body size within populations and assemblages. Science, 381(6662), 1067–1071. 10.1126/science.adg6006

Matta, M. E., & Kimura, D. K. (Eds.). (2012). Age determination manual of the Alaska Fisheries Science Center Age and Growth Program. NOAA Professional Papers NMFS 13:106.

Morrongiello, J. R., Sweetman, P. C., & Thresher, R. E. (2019). Fishing constrains phenotypic responses of marine fish to climate variability. Journal of Animal Ecology, 88(11), 1645– 1656. 10.1111/1365-2656.12999

Neuheimer, A. B., & Grønkjær, P. (2012). Climate effects on size-at-age: Growth in warming waters compensates for earlier maturity in an exploited marine fish. Global Change Biology, 18(6), 1812–1822. 10.1111/j.1365-2486.2012.02673.x

Ohlberger, J. (2013). Climate warming and ectotherm body size—From individual physiology to community ecology. Functional Ecology, 27(4), 991–1001. 10.1111/1365-2435.12098

Ohlberger, J., Cline, T. J., Schindler, D. E., & Lewis, B. (2023). Declines in body size of sockeye salmon associated with increased competition in the ocean. Proceedings of the Royal Society B: Biological Sciences, 290(1992), 20222248. 10.1098/rspb.2022.2248

Ohlberger, J., Ward, E. J., Schindler, D. E., & Lewis, B. (2018). Demographic changes in Chinook salmon across the Northeast Pacific Ocean. Fish and Fisheries, 19(3), 533–546. 10.1111/faf.12272

Oke, K. B., Cunningham, C. J., Westley, P. A. H., Baskett, M. L., Carlson, S. M., Clark, J., Hendry, A. P., Karatayev, V. A., Kendall, N. W., Kibele, J., Kindsvater, H. K., Kobayashi, K. M., Lewis, B., Munch, S., Reynolds, J. D., Vick, G. K., & Palkovacs, E. P. (2020). Recent declines in salmon body size impact ecosystems and fisheries. Nature Communications, 11(1), 4155. 10.1038/s41467-020-17726-z

Oke, K. B., Mueter, F. J., & Litzow, M. A. (2022). Warming leads to opposite patterns in weight-at-age for young versus old age classes of Bering Sea walleye pollock. Canadian Journal of Fisheries and Aquatic Sciences, cjfas-2021–0315. 10.1139/cjfas-2021-0315

Pauly, D. (2021). The gill-oxygen limitation theory (GOLT) and its critics. Science Advances, 7(2), eabc6050. 10.1126/sciadv.abc6050

Peters, R. H. (1986). The ecological implications of body size. Cambridge University Press.

Pilcher, D. J., Naiman, D. M., Cross, J. N., Hermann, A. J., Siedlecki, S. A., Gibson, G. A., & Mathis, J. T. (2019). Modeled Effect of Coastal Biogeochemical Processes, Climate Variability, and Ocean Acidification on Aragonite Saturation State in the Bering Sea. Frontiers in Marine Science, 5, 508. 10.3389/fmars.2018.00508

Pörtner, H.-O. (2010). Oxygen-and capacity-limitation of thermal tolerance: A matrix for integrating climate-related stressor effects in marine ecosystems. Journal of Experimental Biology, 213(6), 881–893. 10.1242/jeb.037523

Rohan, S. K., & O’Leary, C. A. (2024). akfishcondition: Groundfish morphometric condition indicator (Version 4.1.3.).

Rollinson, N., & Rowe, L. (2018). Temperature-dependent oxygen limitation and the rise of Bergmann’s rule in species with aquatic respiration: Temperature-dependent oxygen limitation. Evolution, 72(4), 977–988. 10.1111/evo.13458

Rubalcaba, J. G., Verberk, W. C. E. P., Hendriks, A. J., Saris, B., & Woods, H. A. (2020). Oxygen limitation may affect the temperature and size dependence of metabolism in aquatic ectotherms. Proceedings of the National Academy of Sciences, 117(50), 31963– 31968. 10.1073/pnas.2003292117

Schulte, P. M., Healy, T. M., & Fangue, N. A. (2011). Thermal Performance Curves, Phenotypic Plasticity, and the Time Scales of Temperature Exposure. Integrative and Comparative Biology, 51(5), 691–702. 10.1093/icb/icr097

Sheridan, J. A., & Bickford, D. (2011). Shrinking body size as an ecological response to climate change. Nature Climate Change, 1(8), 401–406. 10.1038/nclimate1259

Solokas, M. A., Feiner, Z. S., Al-Chokachy, R., Budy, P., DeWeber, J. T., Sarvala, J., Sass, G. G., Tolentino, S. A., Walsworth, T. E., & Jensen, O. P. (2023). Shrinking body size and climate warming: Many freshwater salmonids do not follow the rule. Global Change Biology, 29(9), 2478–2492. 10.1111/gcb.16626

Stawitz, C. C., Essington, T. E., Branch, T. A., Haltuch, M. A., Hollowed, A. B., & Spencer, P. D. (2015). A state-space approach for detecting growth variation and application to North Pacific groundfish. Canadian Journal of Fisheries and Aquatic Sciences, 72(9), 1316– 1328. 10.1139/cjfas-2014-0558

Thorson, J. T. (2019). Guidance for decisions using the Vector Autoregressive Spatio-Temporal (VAST) package in stock, ecosystem, habitat and climate assessments. Fisheries Research, 210, 143–161. 10.1016/j.fishres.2018.10.013

Thorson, J. T., Anderson, S. C., Goddard, P., & Rooper, C. (2024). tinyVAST: R package with an expressive interface to specify lagged and simultaneous effects in multivariate spatio-temporal models. 10.48550/arXiv.2401.10193

Thorson, J. T., Barnes, C. L., Friedman, S. T., Morano, J. L., & Siple, M. C. (2023). Spatially varying coefficients can improve parsimony and descriptive power for species distribution models. Ecography, 2023(5), e06510. 10.1111/ecog.06510

Thorson, J. T., & Barnett, L. A. K. (2017). Comparing estimates of abundance trends and distribution shifts using single- and multispecies models of fishes and biogenic habitat. ICES Journal of Marine Science, 74(5), 1311–1321. 10.1093/icesjms/fsw193

Thorson, J. T., & Kristensen, K. (2024). Spatio-temporal Models for Ecologists. CRC Press.

Thorson, J. T., Monnahan, C. C., & Cope, J. M. (2015). The potential impact of time-variation in vital rates on fisheries management targets for marine fishes. Fisheries Research, 169, 8–17. 10.1016/j.fishres.2015.04.007

Van Denderen, D., Gislason, H., Van Den Heuvel, J., & Andersen, K. H. (2020). Global analysis of fish growth rates shows weaker responses to temperature than metabolic predictions. Global Ecology and Biogeography, 29(12), 2203–2213. 10.1111/geb.13189

Van Rijn, I., Buba, Y., DeLong, J., Kiflawi, M., & Belmaker, J. (2017). Large but uneven reduction in fish size across species in relation to changing sea temperatures. Global Change Biology, 23(9), 3667–3674. 10.1111/gcb.13688

Verberk, W. C. E. P., Atkinson, D., Hoefnagel, K. N., Hirst, A. G., Horne, C. R., & Siepel, H. (2021). Shrinking body sizes in response to warming: Explanations for the temperature– size rule with special emphasis on the role of oxygen. Biological Reviews, 96(1), 247–268. 10.1111/brv.12653

Waples, R. S., & Audzijonyte, A. (2016). Fishery-induced evolution provides insights into adaptive responses of marine species to climate change. Frontiers in Ecology and the Environment, 14(4), 217–224. 10.1002/fee.1264

Ward, E. J., Barnett, L. A. K., Anderson, S. C., Commander, C. J. C., & Essington, T. E. (2022). Incorporating non-stationary spatial variability into dynamic species distribution models. ICES Journal of Marine Science, 79(9), 2422–2429. 10.1093/icesjms/fsac179

Wong, S., Bigman, J. S., & Dulvy, N. K. (2021). The metabolic pace of life histories across fishes. Proceedings of the Royal Society B: Biological Sciences, 288(20210910).

