## Supplementary Material for "Changes in body size with age do not follow the temperature-size rule"

This file contains:
Supplementary Tables
Supplementary Figures

Supplementary Tables

Table S1. Number of samples and range of collection years per species.

| Species | Year range | Sample size |
|---|---|---|
| arrowtooth flounder | 1996-2021 | 6021 |
| Pacific cod | 1993-2022 | 32161 |
| walleye pollock | 1991-2023 | 40930 |
| yellowfin sole | 1987-2022 | 20222 |

**Table S2. Models with age-specific random fields are more supported than models with shared spatial and spatiotemporal fields across ages for arrowtooth flounder**. Comparison of models fitted with restricted maximum likelihood (REML) based on marginal AIC for arrowtooth flounder. Models include those with different functional forms (second column), different covariates (temperature, oxygen, none), an interaction between temperature or oxygen and age class (allowing the relationship to vary by age class, e.g., age-specific effects), and different random field structures (models with independent spatial and spatiotemporal effects [‘age-specific effects’], models with shared spatial and spatiotemporal effects across ages [‘shared spatial/spatiotemporal fields’], and models with no spatiotemporal effect and a shared spatial effect across ages [‘shared spatial field’]). For this comparison (using REML), only random effect structures are to be compared (i.e., last three columns, ‘age-specific effects’, ‘shared spatial/spatiotemporal fields’, and ‘shared spatial field’. The bolded values indicate the lowest AIC values (and most supported random effect structure).

| | | | | AIC values | | |
|---|---|---|---|---|---|---|
| Species | Functional form | Covariate | Interaction? | age-specific fields | shared spatial/spatiotemporal fields | shared spatial field |
| arrowtooth flounder | no covariate | no covariate | no | **-5036.85** | -4704.65 | -4173.74 |
| arrowtooth flounder | linear | temperature | no | **-5029.21** | -4696.55 | -4164.95 |
| arrowtooth flounder | linear | oxygen | no | **-5026.59** | -4694.79 | -4164.13 |
| arrowtooth flounder | 2nd order polynomial | temperature | no | **-5026.30** | -4687.21 | -4156.41 |
| arrowtooth flounder | 2nd order polynomial | oxygen | no | **-5018.46** | -4687.81 | -4164.13 |
| arrowtooth flounder | 3rd order polynomial | temperature | no | **-5017.06** | -4677.79 | -4154.22 |
| arrowtooth flounder | 3rd order polynomial | oxygen | no | **-5009.48** | -4679.15 | -4156.51 |
| arrowtooth flounder | linear | oxygen | yes | **-4966.14** | -4633.29 | -4102.03 |
| arrowtooth flounder | linear | temperature | yes | **-4961.29** | -4653.19 | -4117.55 |
| arrowtooth flounder | 2nd order polynomial | oxygen | yes | **-4911.60** | -4593.95 | -4064.97 |
| arrowtooth flounder | 2nd order polynomial | temperature | yes | **-4892.95** | -4578.60 | -4043.39 |
| arrowtooth flounder | 3rd order polynomial | oxygen | yes | **-4860.93** | -4557.68 | -4024.25 |
| arrowtooth flounder | 3rd order polynomial | temperature | yes | **-4816.86** | -4516.35 | -4001.59 |

**Table S3. Same as Table S2 but for Pacific Cod.** Models with age-specific random fields are more supported than models with shared spatial and spatiotemporal fields across ages for Pacific cod**.** Models include those with different functional forms (second column), different covariates (temperature, oxygen, none), an interaction between temperature or oxygen and age class (allowing the relationship to vary by age class, e.g., age-specific effects), and different random field structures (models with independent spatial and spatiotemporal effects ['age-specific effects'], models with shared spatial and spatiotemporal effects across ages ['shared spatial and spatiotemporal fields'], and models with no spatiotemporal effect and a shared spatial effect across ages ['shared spatial field']). For this comparison (using REML), only random effect structures are to be compared (i.e., last three columns, 'age-specific effects', 'shared spatial and spatiotemporal fields', and 'shared spatial field'. The bolded values indicate the lowest AIC values (and most supported random effect structure).

| | | | | AIC values | | |
|---|---|---|---|---|---|---|
| Species | Functional form | Covariate | Interaction? | age-specific fields | shared spatial/spatiotemporal fields | shared spatial field |
| Pacific cod | no covariate | no covariate | no | **-27445.92** | -20372.85 | -12007.10 |
| Pacific cod | linear | temperature | no | **-27438.53** | -20362.79 | -12490.78 |
| Pacific cod | linear | oxygen | no | **-27436.00** | -20361.93 | -12008.99 |
| Pacific cod | 2nd order polynomial | temperature | no | **-27428.65** | -20351.84 | -12479.51 |
| Pacific cod | 2nd order polynomial | oxygen | no | **-27427.37** | -20353.51 | -12142.43 |
| Pacific cod | 3rd order polynomial | oxygen | no | **-27422.05** | -20344.33 | -12247.58 |
| Pacific cod | 3rd order polynomial | temperature | no | **-27415.79** | -20341.53 | -12533.43 |
| Pacific cod | linear | temperature | yes | **-27377.87** | -20401.21 | -12522.97 |
| Pacific cod | linear | oxygen | yes | **-27371.56** | -20309.67 | -11948.36 |
| Pacific cod | 2nd order polynomial | oxygen | yes | **-27313.45** | -20448.16 | -12195.42 |
| Pacific cod | 2nd order polynomial | temperature | yes | **-27307.55** | -20354.24 | -12478.01 |
| Pacific cod | 3rd order polynomial | oxygen | yes | **-27262.29** | -20502.77 | -12354.32 |
| Pacific cod | 3rd order polynomial | temperature | yes | **-27215.18** | -20283.12 | -12441.73 |

**Table S4. Same as Table S2 but for walleye pollock.** Models with age-specific random fields are more supported than models with shared spatial and spatiotemporal fields across ages for walleye pollock. Models include those with different functional forms (second column), different covariates (temperature, oxygen, none), an interaction between temperature or oxygen and age class (allowing the relationship to vary by age class, e.g., age-specific effects), and different random field structures (models with independent spatial and spatiotemporal effects [‘age-specific effects’], models with shared spatial and spatiotemporal effects across ages [‘shared spatial and spatiotemporal fields’], and models with no spatiotemporal effect and a shared spatial effect across ages [‘shared spatial field’]). For this comparison (using REML), only random effect structures are to be compared (i.e., last three columns, ‘age-specific effects’, ‘shared spatial and spatiotemporal fields’, and ‘shared spatial field’.

| | | | | AIC values | | |
|---|---|---|---|---|---|---|
| Species | Functional form | Covariate | Interaction? | age-specific fields | shared spatial/spatiotemporal fields | shared spatial field |
| walleye pollock | 2nd order polynomial | temperature | no | **-58709.12** | -41627.85 | -36382.62 |
| walleye pollock | linear | temperature | no | **-58697.66** | -41639.97 | -36385.56 |
| walleye pollock | 3rd order polynomial | temperature | no | **-58695.13** | -41618.58 | -36407.09 |
| walleye pollock | no covariate | no covariate | no | **-58694.35** | -41651.12 | -36395.88 |
| walleye pollock | linear | oxygen | no | **-58682.40** | -41639.43 | -36384.46 |
| walleye pollock | 2nd order polynomial | oxygen | no | **-58671.78** | -41629.81 | -36380.86 |
| walleye pollock | 3rd order polynomial | oxygen | no | **-58670.84** | -41619.80 | -36379.25 |
| walleye pollock | linear | temperature | yes | **-58560.63** | -42646.58 | -37639.16 |
| walleye pollock | linear | oxygen | yes | **-58530.25** | -41774.97 | -36575.66 |
| walleye pollock | 2nd order polynomial | temperature | yes | **-58413.94** | -42613.51 | -37669.12 |
| walleye pollock | 2nd order polynomial | oxygen | yes | **-58412.67** | -42221.18 | -37271.75 |
| walleye pollock | 3rd order polynomial | oxygen | yes | **-58292.96** | -42382.86 | -37453.54 |
| walleye pollock | 3rd order polynomial | temperature | yes | **-58221.64** | -42454.51 | -37515.05 |

**Table S5. Same as Table S2 but for yellowfin sole.** Models with age-specific random fields are more supported than models with shared spatial and spatiotemporal fields across ages for yellowfin sole. Models include those with different functional forms (second column), different covariates (temperature, oxygen, none), an interaction between temperature or oxygen and age class (allowing the relationship to vary by age class, e.g., age-specific effects), and different random field structures (models with independent spatial and spatiotemporal effects [‘age-specific effects’], models with shared spatial and spatiotemporal effects across ages [‘shared spatial and spatiotemporal fields’], and models with no spatiotemporal effect and a shared spatial effect across ages [‘shared spatial field’]). For this comparison (using REML), only random effect structures are to be compared (i.e., last three columns, ‘age-specific effects’, ‘shared spatial and spatiotemporal fields’, and ‘shared spatial field’. The bolded values indicate the lowest AIC values (and most supported random effect structure).

| | | | | AIC values | | |
|---|---|---|---|---|---|---|
| Species | Functional form | Covariate | Interaction? | age-specific fields | shared spatial/spatiotemporal fields | shared spatial field |
| yellowfin sole | no covariate | no covariate | no | **-21376.46** | -17057.76 | -13204.08 |
| yellowfin sole | linear | temperature | no | **-21371.54** | -17048.46 | -13272.10 |
| yellowfin sole | linear | oxygen | no | **-21369.19** | -17047.75 | -13200.41 |
| yellowfin sole | 2nd order polynomial | temperature | no | **-21362.41** | -17038.71 | -13260.49 |
| yellowfin sole | 2nd order polynomial | oxygen | no | **-21359.05** | -17041.30 | -13191.18 |
| yellowfin sole | 3rd order polynomial | oxygen | no | **-21349.75** | -17033.70 | -13200.48 |
| yellowfin sole | 3rd order polynomial | temperature | no | **-21349.58** | -17027.83 | -13301.82 |
| yellowfin sole | linear | temperature | yes | **-21189.45** | -17021.30 | -13245.63 |
| yellowfin sole | linear | oxygen | yes | **-21180.06** | -16876.15 | -13039.18 |
| yellowfin sole | 2nd order polynomial | oxygen | yes | **-21018.98** | -16754.20 | -12919.25 |
| yellowfin sole | 2nd order polynomial | temperature | yes | **-20968.12** | -16898.53 | -13090.94 |
| yellowfin sole | 3rd order polynomial | oxygen | yes | **-20858.99** | -16736.34 | -12881.19 |
| yellowfin sole | 3rd order polynomial | temperature | yes | **-20720.31** | -16713.82 | -12946.45 |

**Table S6. Models with temperature and no interaction between temperature and age are the most supported models explaining changes in weight-at-age over time for arrowtooth flounder.** Comparison of models fitted with maximum likelihood (ML) based on marginal AIC for arrowtooth flounder. Models include those with different functional forms (second column), different covariates (temperature, oxygen, none), and an interaction between temperature or oxygen and age class (allowing the relationship to vary by age class, e.g., age-specific effects). The random effect structure for all models in this comparison follow the most supported structure, age-specific spatial and spatiotemporal fields. For this comparison (using ML), only the importance of fixed effects are to be compared (i.e., functional form, covariate, interaction). All models within 2 AIC units of the most supported model are highlighted in grey and the most supported model (that with the lowest AIC) is emboldened.

| Species | Functional form | Covariate | Interaction? | AIC (age-specific fields model) |
|---|---|---|---|---|
| **arrowtooth flounder** | **3rd order polynomial** | **temperature** | **no** | **-5074.870** |
| arrowtooth flounder | 2nd order polynomial | temperature | no | -5074.353 |
| arrowtooth flounder | linear | temperature | no | -5069.105 |
| arrowtooth flounder | no covariate | no covariate | no | -5068.391 |
| arrowtooth flounder | linear | oxygen | no | -5066.520 |
| arrowtooth flounder | 2nd order polynomial | oxygen | no | -5065.642 |
| arrowtooth flounder | 3rd order polynomial | oxygen | no | -5063.828 |
| arrowtooth flounder | linear | oxygen | yes | -5060.757 |
| arrowtooth flounder | linear | temperature | yes | -5055.490 |
| arrowtooth flounder | 2nd order polynomial | oxygen | yes | -5055.454 |
| arrowtooth flounder | 3rd order polynomial | oxygen | yes | -5054.467 |
| arrowtooth flounder | 2nd order polynomial | temperature | yes | -5048.883 |
| arrowtooth flounder | 3rd order polynomial | temperature | yes | -5043.535 |

**Table S7. Same as Table S6 but for Pacific cod.** Models with temperature and no interaction between temperature and age are the most supported models explaining changes in weight-at-age over time for Pacific cod. Comparison of models fitted with maximum likelihood (ML) based on marginal AIC for Pacific cod. Models include those with different functional forms (second column), different covariates (temperature, oxygen, none), and an interaction between temperature or oxygen and age class (allowing the relationship to vary by age class, e.g., age-specific effects). The random effect structure for all models in this comparison follow the most supported structure, age-specific spatial and spatiotemporal fields. For this comparison (using ML), only the importance of fixed effects are to be compared (i.e., functional form, covariate, interaction). All models within 2 AIC units of the most supported model are highlighted in grey and the most supported model (that with the lowest AIC) is emboldened.

| Species | Functional form | Covariate | Interaction? | AIC (age-specific fields model) |
|---|---|---|---|---|
| **Pacific cod** | **2nd order polynomial** | **temperature** | **no** | **-27465.92** |
| Pacific cod | 3rd order polynomial | oxygen | no | -27465.87 |
| Pacific cod | linear | temperature | no | -27465.71 |
| Pacific cod | 2nd order polynomial | temperature | yes | -27464.39 |
| Pacific cod | 3rd order polynomial | temperature | no | -27464.14 |
| Pacific cod | no covariate | no covariate | no | -27463.72 |
| Pacific cod | linear | oxygen | no | -27463.44 |
| Pacific cod | 2nd order polynomial | oxygen | no | -27463.00 |
| Pacific cod | linear | temperature | yes | -27460.73 |
| Pacific cod | linear | oxygen | yes | -27459.15 |
| Pacific cod | 3rd order polynomial | oxygen | yes | -27456.62 |
| Pacific cod | 2nd order polynomial | oxygen | yes | -27454.45 |
| Pacific cod | 3rd order polynomial | temperature | yes | -27453.85 |

**Table S8. Same as Table S6 but for walleye pollock.** Models with temperature and no interaction between temperature and age are the most supported models explaining changes in weight-at-age over time for walleye pollock. Comparison of models fitted with maximum likelihood (ML) based on marginal AIC for walleye pollock. Models include those with different functional forms (second column), different covariates (temperature, oxygen, none), and an interaction between temperature or oxygen and age class (allowing the relationship to vary by age class, e.g., age-specific effects). The random effect structure for all models in this comparison follow the most supported structure, age-specific spatial and spatiotemporal fields. For this comparison (using ML), only the importance of fixed effects are to be compared (i.e., functional form, covariate, interaction). All models within 2 AIC units of the most supported model are highlighted in grey and the most supported model (that with the lowest AIC) is emboldened.

| Species | Functional form | Covariate | Interaction? | AIC (age-specific fields model) |
|---|---|---|---|---|
| **walleye pollock** | **2nd order polynomial** | **temperature** | **no** | **-58785.48** |
| walleye pollock | 3rd order polynomial | temperature | no | -58783.59 |
| walleye pollock | linear | temperature | no | -58763.47 |
| walleye pollock | 3rd order polynomial | oxygen | no | -58756.88 |
| walleye pollock | 2nd order polynomial | temperature | yes | -58752.36 |
| walleye pollock | no covariate | no covariate | no | -58750.45 |
| walleye pollock | linear | temperature | yes | -58750.06 |
| walleye pollock | linear | oxygen | no | -58749.33 |
| walleye pollock | 2nd order polynomial | oxygen | no | -58747.80 |
| walleye pollock | linear | oxygen | yes | -58729.31 |
| walleye pollock | 3rd order polynomial | temperature | yes | -58728.14 |
| walleye pollock | 2nd order polynomial | oxygen | yes | -58718.20 |
| walleye pollock | 3rd order polynomial | oxygen | yes | -58707.63 |

**Table S9. Same as Table S6 but for yellowfin sole.** Models with temperature and no interaction between temperature and age are the most supported models explaining changes in weight-at-age over time for yellowfin sole. Comparison of models fitted with maximum likelihood (ML) based on marginal AIC for yellowfin sole. Models include those with different functional forms (second column), different covariates (temperature, oxygen, none), and an interaction between temperature or oxygen and age class (allowing the relationship to vary by age class, e.g., age-specific effects). The random effect structure for all models in this comparison follow the most supported structure, age-specific spatial and spatiotemporal fields. For this comparison (using ML), only the importance of fixed effects are to be compared (i.e., functional form, covariate, interaction). All models within 2 AIC units of the most supported model are highlighted in grey and the most supported model (that with the lowest AIC) is emboldened.

| Species | Functional form | Covariate | Interaction? | AIC (age-specific fields model) |
|---|---|---|---|---|
| **yellowfin sole** | **2nd order polynomial** | **temperature** | **no** | **-21425.97** |
| yellowfin sole | linear | temperature | no | -21425.27 |
| yellowfin sole | linear | oxygen | no | -21424.76 |
| yellowfin sole | 3rd order polynomial | temperature | no | -21423.98 |
| yellowfin sole | 2nd order polynomial | oxygen | no | -21422.82 |
| yellowfin sole | no covariate | no covariate | no | -21422.12 |
| yellowfin sole | 3rd order polynomial | oxygen | no | -21422.10 |
| yellowfin sole | linear | oxygen | yes | -21397.19 |
| yellowfin sole | linear | temperature | yes | -21391.56 |
| yellowfin sole | 2nd order polynomial | oxygen | yes | -21365.81 |
| yellowfin sole | 2nd order polynomial | temperature | yes | -21350.24 |
| yellowfin sole | 3rd order polynomial | oxygen | yes | -21342.03 |
| yellowfin sole | 3rd order polynomial | temperature | yes | -21305.99 |

**Table S10. If models with suboptimal random effect structures (shared spatial/spatiotemporal or shared spatial) had been used for arrowtooth flounder, the interaction between age class and weight would have been important, providing spurious support for the temperature-size rule.** Comparison of all fitted models using maximum likelihood (ML) estimation for arrowtooth flounder based on AIC. Models include those with different functional forms (second column), different covariates (temperature, oxygen, none), an interaction between temperature or oxygen and age class (allowing the relationship to vary by age class, e.g., age-specific effects), and different random field structures (models with independent spatial and spatiotemporal effects [‘age-specific fields’], models with shared spatial and spatiotemporal effects across ages [‘shared spatial/spatiotemporal fields’], and models with no spatiotemporal effect and a shared spatial effect across ages [‘shared spatial field’]). For this comparison (using ML), the point is to highlight that the most supported models for those with shared spatial and spatiotemporal effects (last two columns, [‘shared spatial/spatiotemporal fields’ and ‘shared spatial field’]) include an interaction between age class and temperature, which is a spurious relationship given both spatial and spatiotemporal random fields should be separate by age class (see Table S2). Models are ordered by lowest AIC for the ‘shared spatial/spatiotemporal fields’ column to highlight this.

| | | | | AIC values | | |
|---|---|---|---|---|---|---|
| Species | Functional form | Covariate | Interaction? | age-specific fields | shared spatial/spatiotemporal fields | shared spatial field |
| arrowtooth flounder | 3rd order polynomial | oxygen | yes | -5054.47 | -4781.69 | -4245.67 |
| arrowtooth flounder | linear | temperature | yes | -5055.49 | -4776.50 | -4240.55 |
| arrowtooth flounder | linear | temperature | yes | -5055.49 | -4776.50 | -4240.55 |
| arrowtooth flounder | 3rd order polynomial | temperature | yes | -5043.53 | -4773.67 | -4257.28 |
| arrowtooth flounder | 2nd order polynomial | oxygen | yes | -5055.45 | -4767.64 | -4237.12 |
| arrowtooth flounder | 2nd order polynomial | temperature | yes | -5048.88 | -4766.05 | -4229.87 |
| arrowtooth flounder | no covariate | no covariate | no | -5068.39 | -4763.75 | -4233.76 |
| arrowtooth flounder | linear | temperature | no | -5069.10 | -4763.62 | -4232.80 |
| arrowtooth flounder | 2nd order polynomial | temperature | no | -5074.35 | -4762.26 | -4232.42 |
| arrowtooth flounder | linear | oxygen | no | -5066.52 | -4761.87 | -4231.77 |
| arrowtooth flounder | 3rd order polynomial | temperature | no | -5074.87 | -4761.76 | -4239.41 |
| arrowtooth flounder | 2nd order polynomial | oxygen | no | -5065.64 | -4761.41 | -4237.92 |
| arrowtooth flounder | 3rd order polynomial | oxygen | no | -5063.83 | -4759.43 | -4236.86 |
| arrowtooth flounder | linear | oxygen | yes | -5060.76 | -4756.00 | -4224.43 |

**Table S11. Same as Table S10 for but Pacific cod.** If models with suboptimal random effect structures (shared spatial/spatiotemporal or shared spatial) had been used for Pacific cod. the interaction between age class and weight would have been important, providing spurious support for the temperature-size rule. Comparison of all fitted models using maximum likelihood (ML) estimation for Pacific cod based on AIC. Models include those with different functional forms (second column), different covariates (temperature, oxygen, none), an interaction between temperature or oxygen and age class (allowing the relationship to vary by age class, e.g., age-specific effects), and different random field structures (models with independent spatial and spatiotemporal effects [‘age-specific fields’], models with shared spatial and spatiotemporal effects across ages [‘shared spatial/spatiotemporal fields’], and models with no spatiotemporal effect and a shared spatial effect across ages [‘shared spatial field’]). For this comparison (using ML), the point is to highlight that the most supported models for those with shared spatial and spatiotemporal effects (last two columns, [‘shared spatial/spatiotemporal fields’ and ‘shared spatial field’]) include an interaction between age class and temperature, which is a spurious term given both spatial and spatiotemporal random fields should be separate by age class (see Table S3). Models are ordered by lowest AIC for the ‘shared spatial/spatiotemporal fields’ column to highlight this.

| | | | | AIC values | | |
|---|---|---|---|---|---|---|
| Species | Functional form | Covariate | Interaction? | age-specific fields | shared spatial/spatiotemporal fields | shared spatial field |
| Pacific cod | 3rd order polynomial | oxygen | yes | -27456.62 | -20751.34 | -12595.87 |
| Pacific cod | 2nd order polynomial | oxygen | yes | -27454.45 | -20641.49 | -12383.67 |
| Pacific cod | 3rd order polynomial | temperature | yes | -27453.85 | -20581.28 | -12735.26 |
| Pacific cod | 2nd order polynomial | temperature | yes | -27464.39 | -20568.40 | -12688.61 |
| Pacific cod | linear | temperature | yes | -27460.73 | -20537.58 | -12656.92 |
| Pacific cod | linear | oxygen | yes | -27459.15 | -20446.78 | -12082.16 |
| Pacific cod | no covariate | no covariate | no | -27463.72 | -20438.91 | -12072.21 |
| Pacific cod | linear | temperature | no | -27465.71 | -20437.17 | -12564.65 |
| Pacific cod | linear | oxygen | no | -27463.44 | -20436.98 | -12082.98 |
| Pacific cod | 2nd order polynomial | oxygen | no | -27463.00 | -20435.76 | -12224.03 |
| Pacific cod | 2nd order polynomial | temperature | no | -27465.92 | -20435.42 | -12563.43 |
| Pacific cod | 3rd order polynomial | temperature | no | -27464.14 | -20435.21 | -12628.23 |
| Pacific cod | 3rd order polynomial | oxygen | no | -27465.87 | -20433.82 | -12336.50 |

**Table S12. Same as Table S10 but for walleye pollock.** If models with suboptimal random effect structures (shared spatial/spatiotemporal or shared spatial) had been used for walleye pollock, the interaction between age class and weight would have been important, providing spurious support for the temperature-size rule. Comparison of all fitted models using maximum likelihood (ML) estimation for walleye pollock based on AIC. Models include those with different functional forms (second column), different covariates (temperature, oxygen, none), an interaction between temperature or oxygen and age class (allowing the relationship to vary by age class, e.g., age-specific effects), and different random field structures (models with independent spatial and spatiotemporal effects ['age-specific fields'], models with shared spatial and spatiotemporal effects across ages ['shared spatial/spatiotemporal fields'], and models with no spatiotemporal effect and a shared spatial effect across ages ['shared spatial field']). For this comparison (using ML), the point is to highlight that the most supported models for those with shared spatial and spatiotemporal effects (last two columns, ['shared spatial/spatiotemporal fields' and 'shared spatial field']) include an interaction between age class and temperature, which is a spurious term given both spatial and spatiotemporal random fields should be separate by age class (see Table S4). Models are ordered by lowest AIC for the 'shared spatial/spatiotemporal fields' column to highlight this.

| | | | | AIC values | | |
|---|---|---|---|---|---|---|
| Species | Functional form | Covariate | Interaction? | age-specific fields | shared spatial/spatiotemporal fields | shared spatial field |
| walleye pollock | 3rd order polynomial | temperature | yes | -58728.14 | -43054.45 | -38109.55 |
| walleye pollock | 2nd order polynomial | temperature | yes | -58752.36 | -43043.59 | -38094.62 |
| walleye pollock | linear | temperature | yes | -58750.06 | -42922.17 | -37911.27 |
| walleye pollock | 3rd order polynomial | oxygen | yes | -58707.63 | -42876.07 | -37937.54 |
| walleye pollock | 2nd order polynomial | oxygen | yes | -58718.20 | -42606.08 | -37649.79 |
| walleye pollock | linear | oxygen | yes | -58729.31 | -42050.51 | -36846.44 |
| walleye pollock | no covariate | no covariate | no | -58750.45 | -41786.11 | -36528.67 |
| walleye pollock | linear | oxygen | no | -58749.33 | -41784.23 | -36526.93 |
| walleye pollock | linear | temperature | no | -58763.47 | -41784.21 | -36527.98 |
| walleye pollock | 3rd order polynomial | temperature | no | -58783.59 | -41784.12 | -36572.42 |
| walleye pollock | 2nd order polynomial | oxygen | no | -58747.80 | -41782.62 | -36531.48 |
| walleye pollock | 2nd order polynomial | temperature | no | -58785.48 | -41782.23 | -36535.92 |
| walleye pollock | 3rd order polynomial | oxygen | no | -58756.88 | -41780.72 | -36538.05 |

**Table S13. Same as Table S10 but for yellowfin sole.** If models with suboptimal random effect structures (shared spatial/spatiotemporal or shared spatial) had been used for yellowfin sole, the interaction between age class and weight would have been important, providing spurious support for the temperature-size rule. Comparison of all fitted models using maximum likelihood (ML) estimation for yellowfin sole based on AIC. Models include those with different functional forms (second column), different covariates (temperature, oxygen, none), an interaction between temperature or oxygen and age class (allowing the relationship to vary by age class, e.g., age-specific effects), and different random field structures (models with independent spatial and spatiotemporal effects [‘age-specific fields’], models with shared spatial and spatiotemporal effects across ages [‘shared spatial/spatiotemporal fields’], and models with no spatiotemporal effect and a shared spatial effect across ages [‘shared spatial field’]). For this comparison (using ML), the point is to highlight that the most supported models for those with shared spatial and spatiotemporal effects (last two columns, [‘shared spatial/spatiotemporal fields’ and ‘shared spatial field’]) include an interaction between age class and temperature, which is a spurious term given both spatial and spatiotemporal random fields should be separate by age class (see Table S5). Models are ordered by lowest AIC for the ‘shared spatial/spatiotemporal fields’ column to highlight this.

| | | | | AIC values | | |
|---|---|---|---|---|---|---|
| Species | Functional form | Covariate | Interaction? | age-specific fields | shared spatial/spatiotemporal fields | shared spatial field |
| yellowfin sole | 3rd order polynomial | temperature | yes | -21305.99 | -17431.49 | -13646.38 |
| yellowfin sole | 2nd order polynomial | temperature | yes | -21350.24 | -17415.63 | -13594.16 |
| yellowfin sole | linear | temperature | yes | -21391.56 | -17353.78 | -13568.37 |
| yellowfin sole | 3rd order polynomial | oxygen | yes | -21342.03 | -17328.66 | -13452.88 |
| yellowfin sole | no covariate | no covariate | no | -21422.12 | -17221.95 | -13362.92 |
| yellowfin sole | 2nd order polynomial | oxygen | no | -21422.82 | -17220.59 | -13365.79 |
| yellowfin sole | linear | oxygen | no | -21424.76 | -17220.27 | -13367.77 |
| yellowfin sole | linear | temperature | no | -21425.27 | -17220.12 | -13439.22 |
| yellowfin sole | 3rd order polynomial | oxygen | no | -21422.10 | -17220.04 | -13382.67 |
| yellowfin sole | 2nd order polynomial | temperature | no | -21425.97 | -17218.70 | -13437.22 |
| yellowfin sole | 3rd order polynomial | temperature | no | -21423.98 | -17217.04 | -13488.94 |
| yellowfin sole | 2nd order polynomial | oxygen | yes | -21365.81 | -17214.33 | -13363.83 |
| yellowfin sole | linear | oxygen | yes | -21397.19 | -17208.55 | -13360.96 |

Supplementary Figures

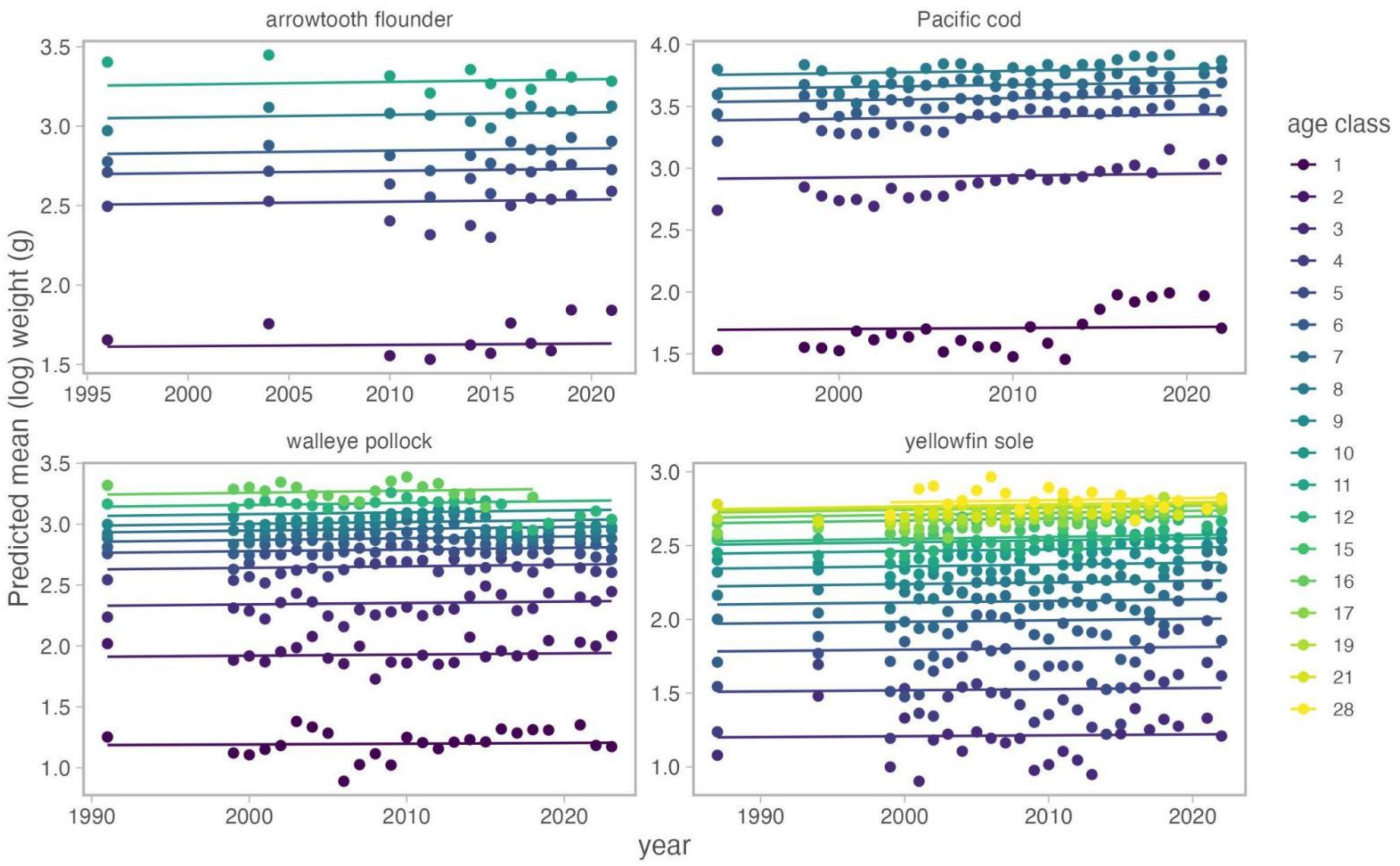

Figure S1. There was no long-term change in weight-at-age over time for any of the four species examined in this study. Each panel corresponds to a species (name on top of panel). Prediction lines come from spatiotemporal models for each age of each species fit to weight data with a year (continuous) fixed effect and spatial and spatiotemporal random fields.

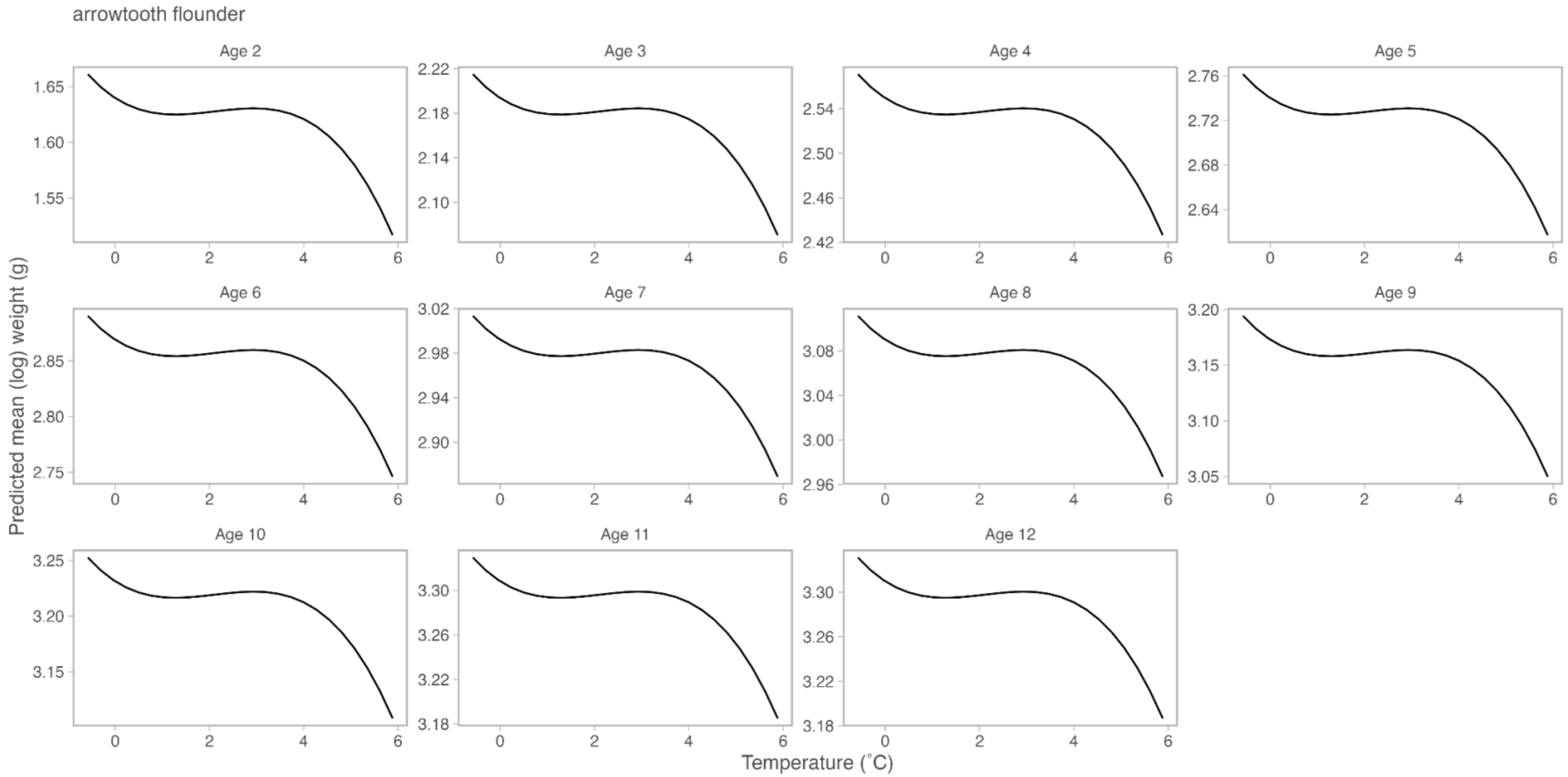

Figure S2. The relationship between weight and temperature by age for arrowtooth flounder. Each panel shows the relationship between temperature for one age class (labeled on top of panel). Fit lines show the predicted weight at a given temperature value from the most supported model with (that with age-specific fields, see text and Table S6).

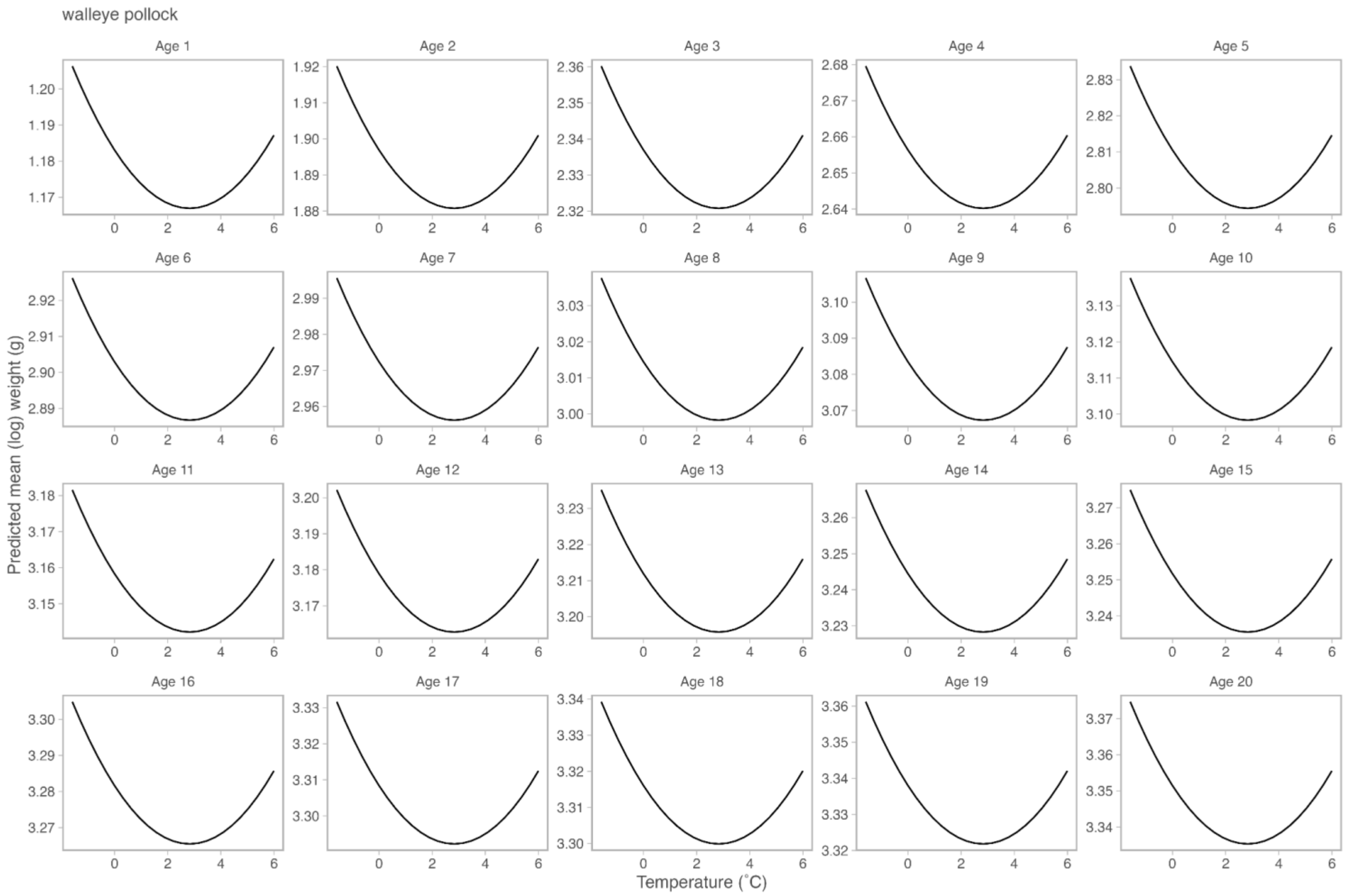

Figure S3. The relationship between weight and temperature by age for walleye pollock. Each panel shows the relationship between temperature for one age class (labeled on top of panel). Fit lines show the predicted weight at a given temperature value from the most supported model with (that with age-specific fields, see text and Table S7).

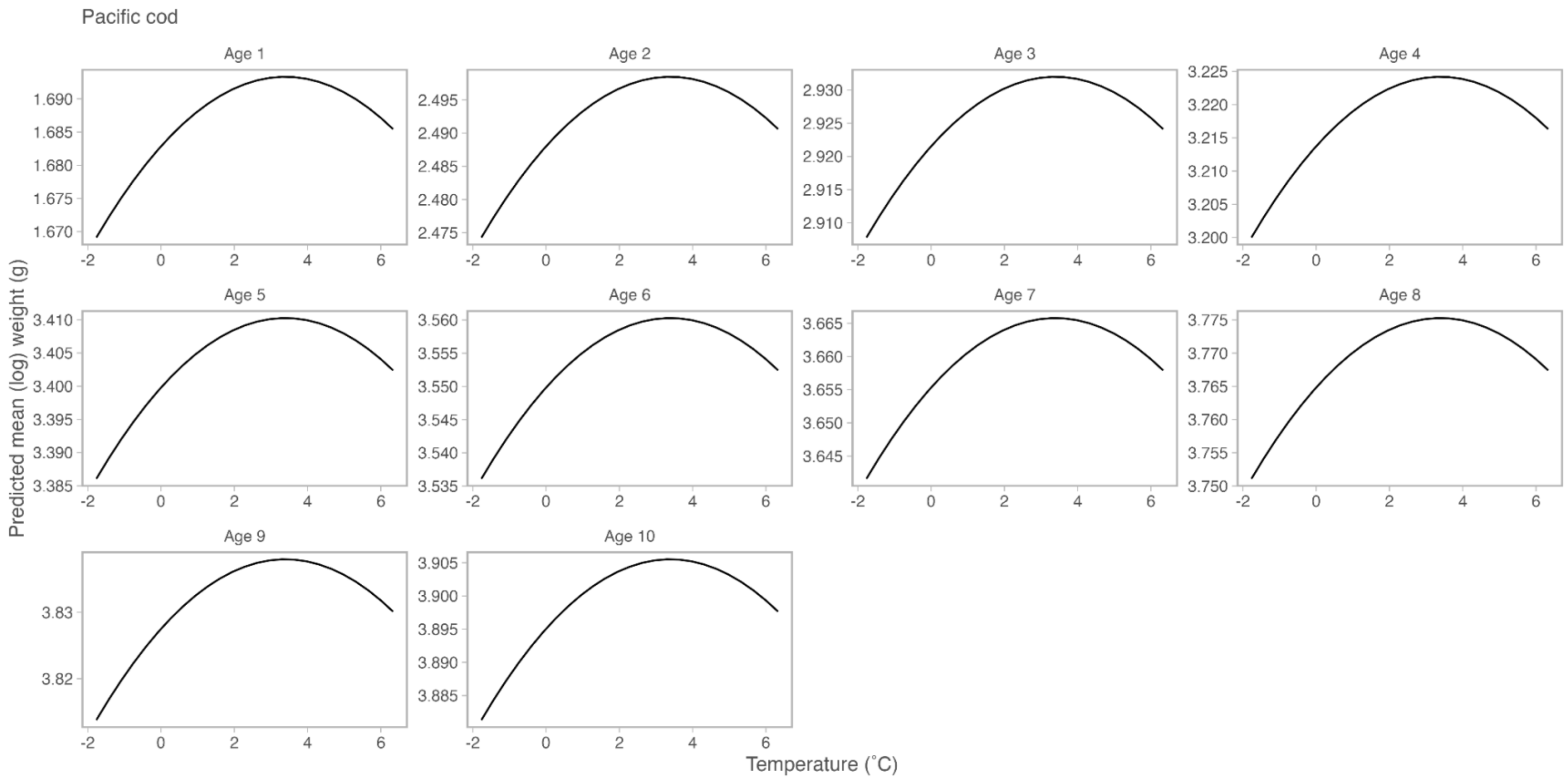

Figure S4. The relationship between weight and temperature by age for Pacific cod. Each panel shows the relationship between temperature for one age class (labeled on top of panel). Fit lines show the predicted weight at a given temperature value from the most supported model with (that with age-specific fields, see text and Table S8).

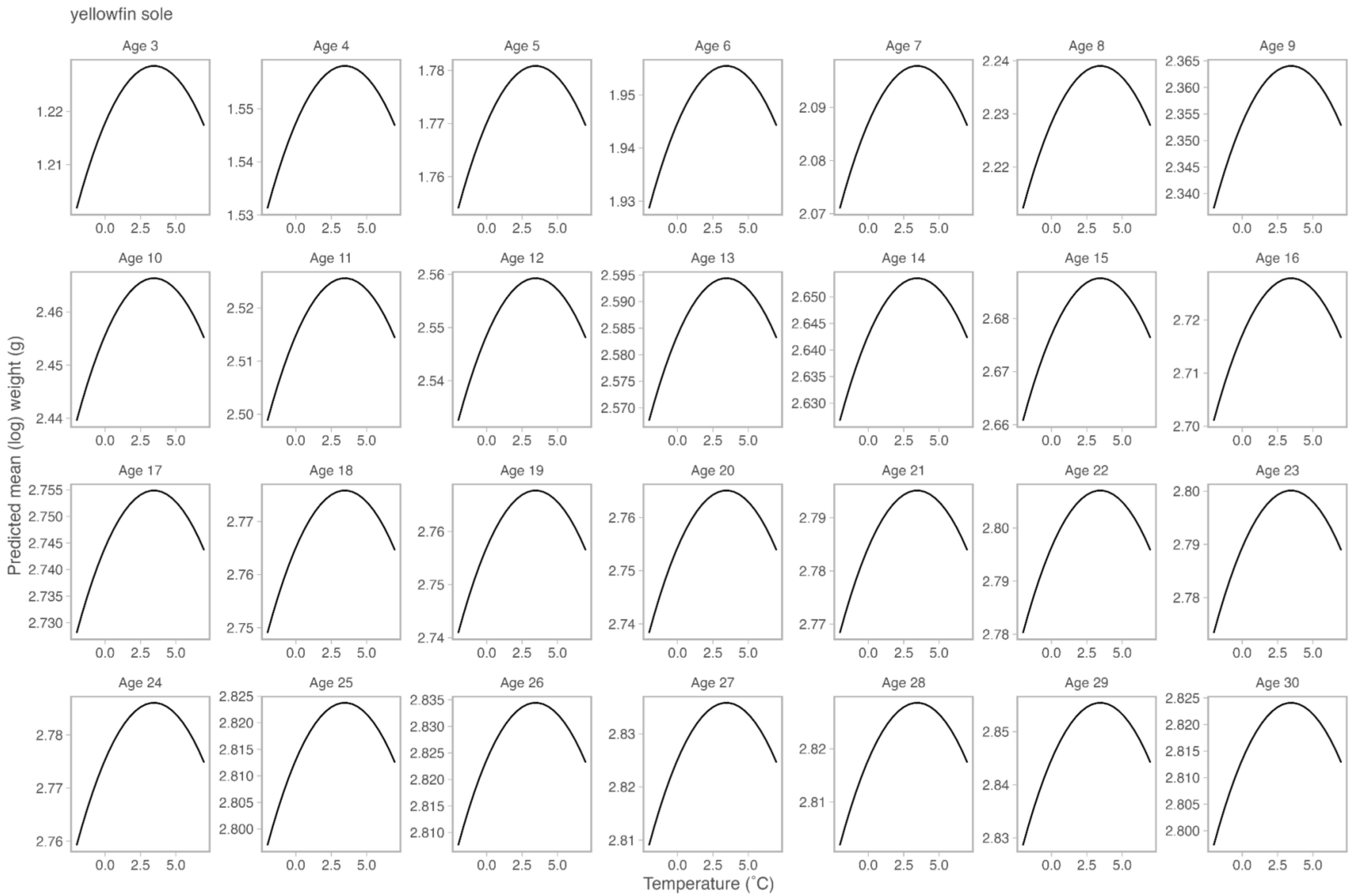

Figure S5. The relationship between weight and temperature by age for yellowfin sole. Each panel shows the relationship between temperature for one age class (labeled on top of panel). Fit lines show the predicted weight at a given temperature value from the most supported model with (that with age-specific fields, see text and Table S9).

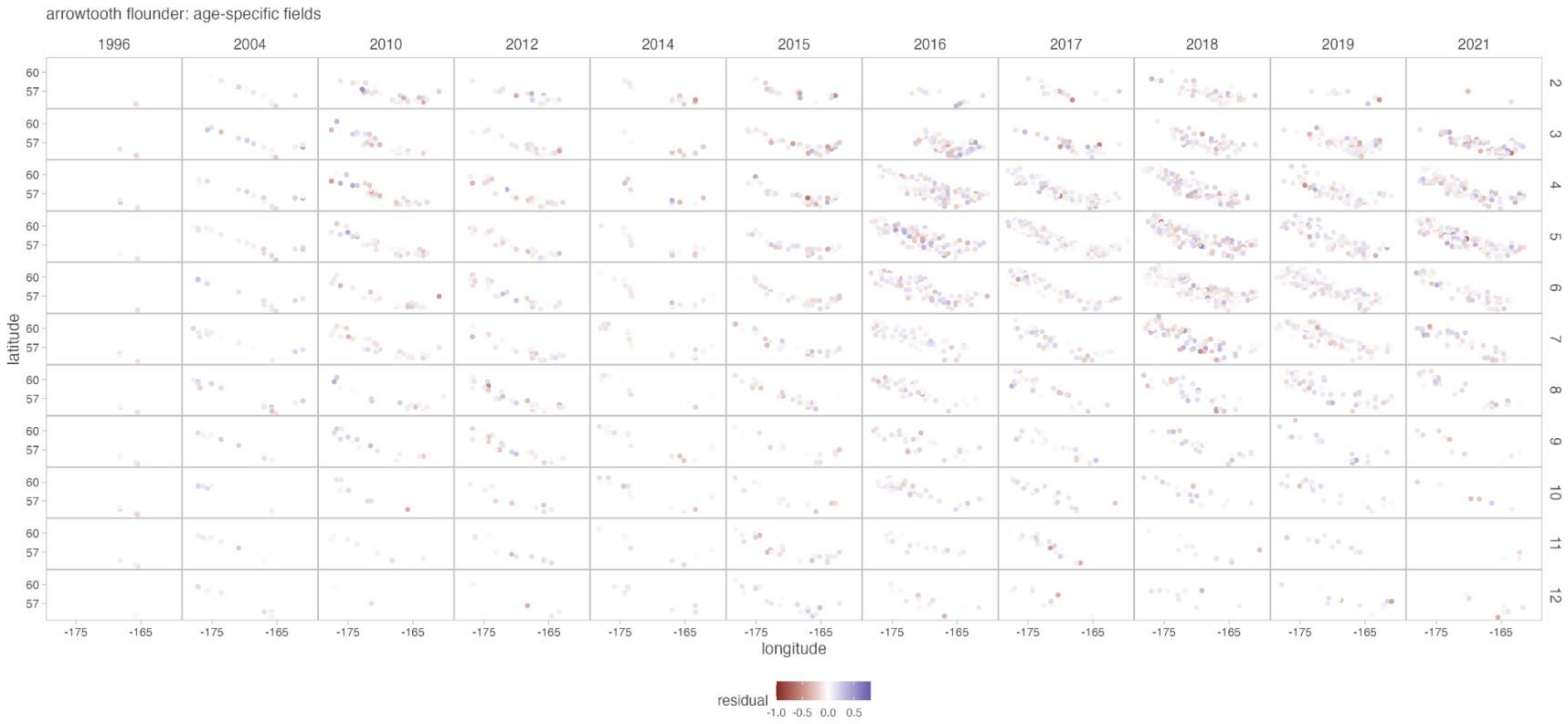

arrowtooth flounder: age-specific fields
1996
2004
2010
2012
2014
2015
2016
2017
2018
2019
2021
60
57
latitude
-175
-165
longitude
residual
-1.0
-0.5
0.0
0.5

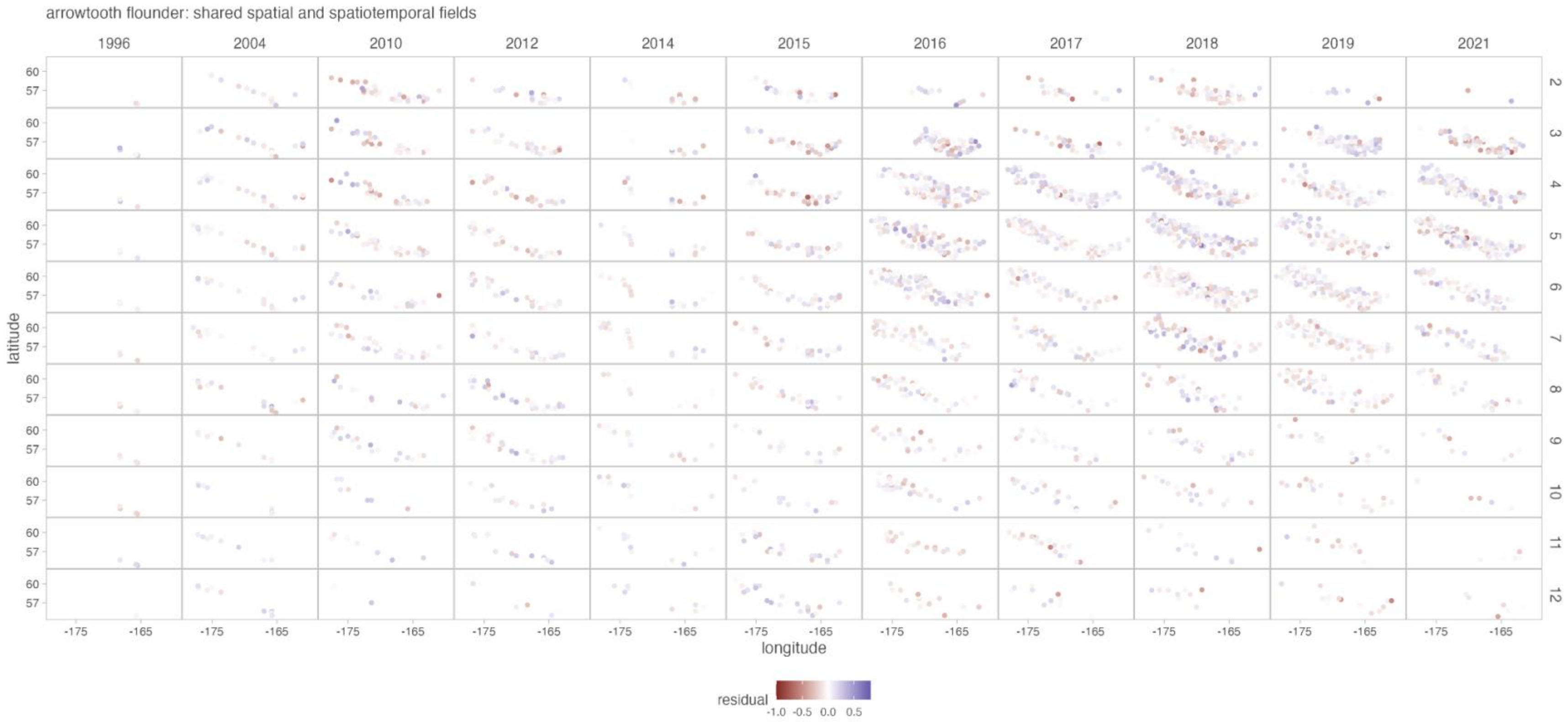

arrowtooth flounder: shared spatial and spatiotemporal fields
1996
2004
2010
2012
2014
2015
2016
2017
2018
2019
2021
60
57
latitude
-175
-165
longitude
residual
-1.0
-0.5
0.0
0.5

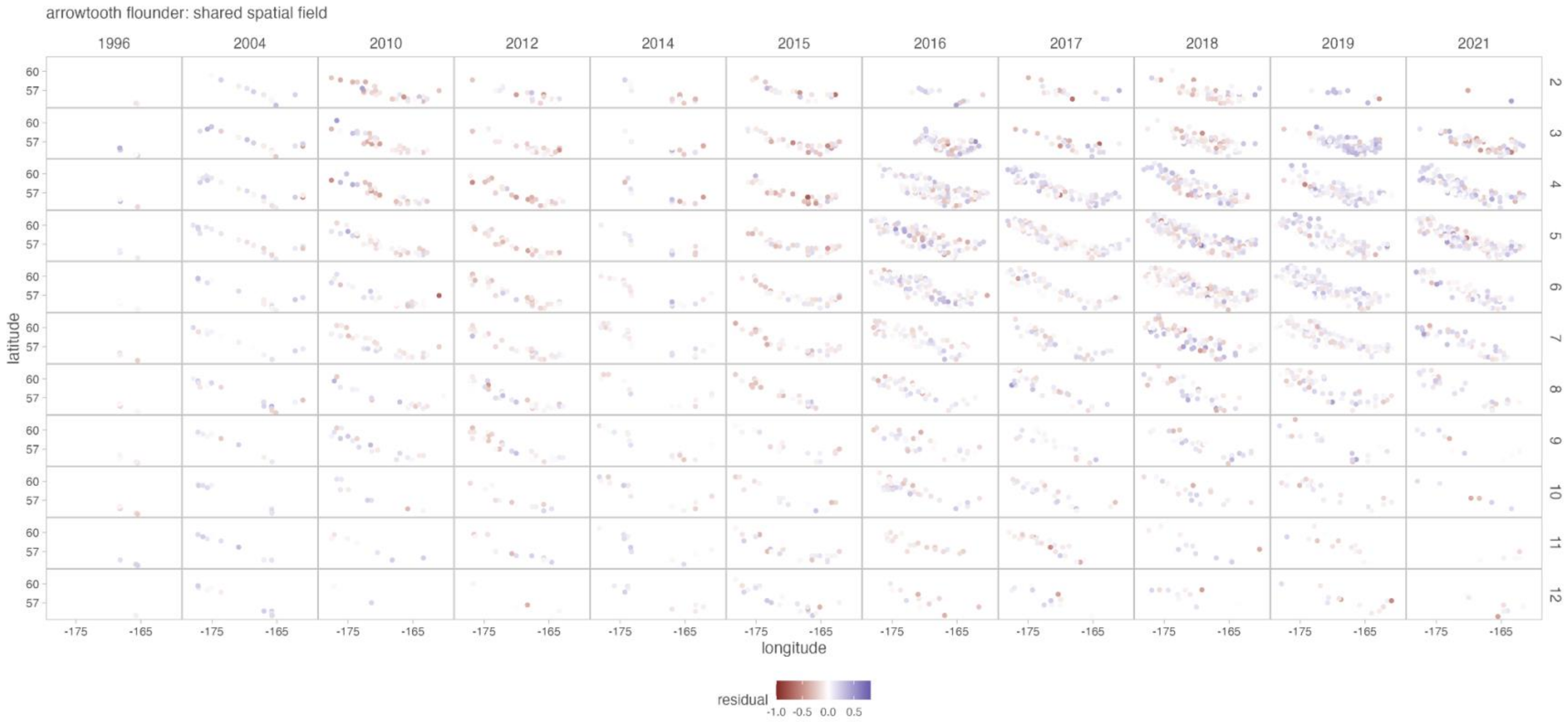

Figure S6. Comparison of residuals over space and time for arrowtooth flounder among models with the same fixed effect structure but the three different random effect structures (see text): age-specific fields (first figure), shared spatial and spatiotemporal fields (second figure), and shared spatial fields (third figure). Residuals shown here are from the most supported model with the most supported random effect structure, i.e., the most supported model in Table S6, which has a third order polynomial relationship between weight and temperature with no interaction between age and temperature. Age class is indicated by row (i.e., 2 = Age 2).

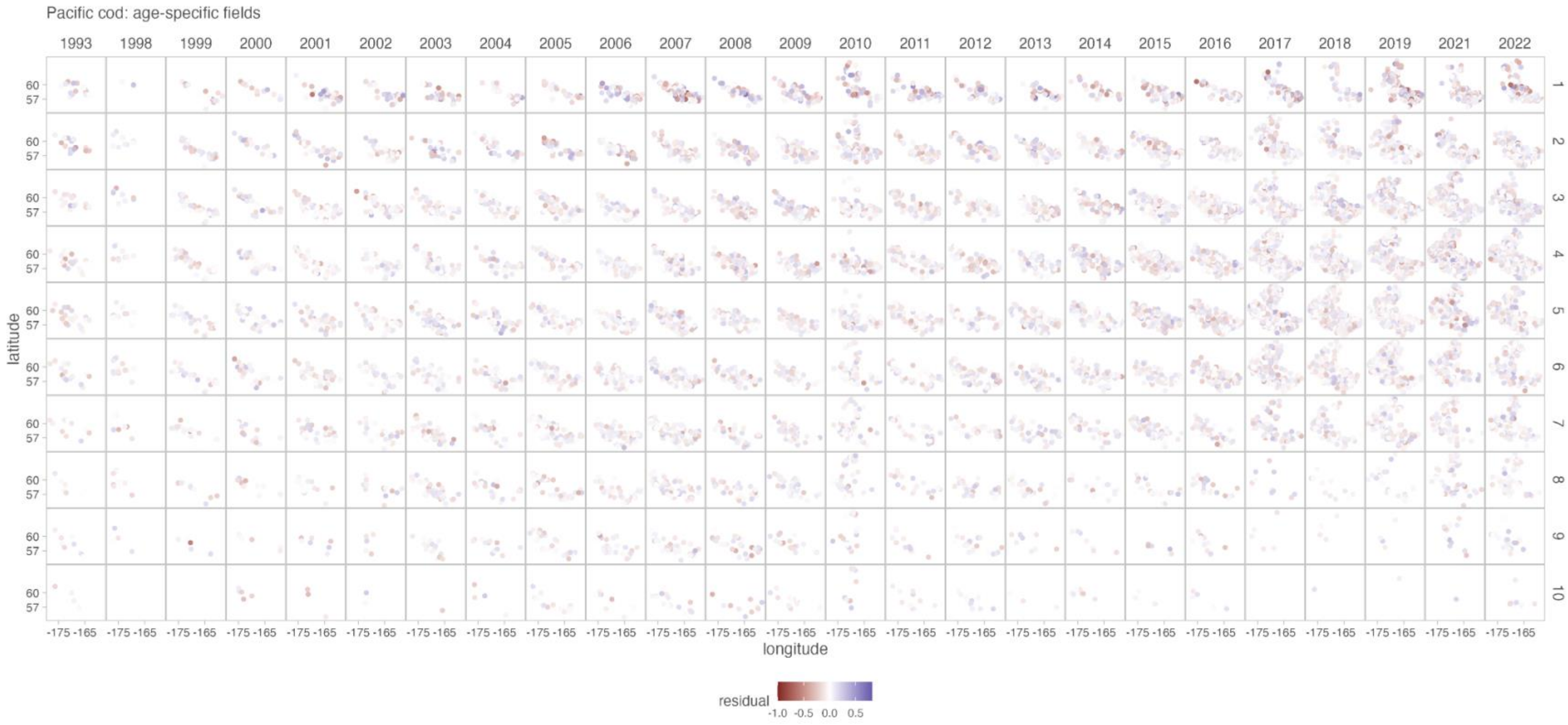

Pacific cod: age-specific fields
1993
1998
1999
2000
2001
2002
2003
2004
2005
2006
2007
2008
2009
2010
2011
2012
2013
2014
2015
2016
2017
2018
2019
2021
2022
1
2
3
4
5
6
7
8
9
10
latitude
60
57
longitude
-175 -165
residual
-1.0 -0.5 0.0 0.5

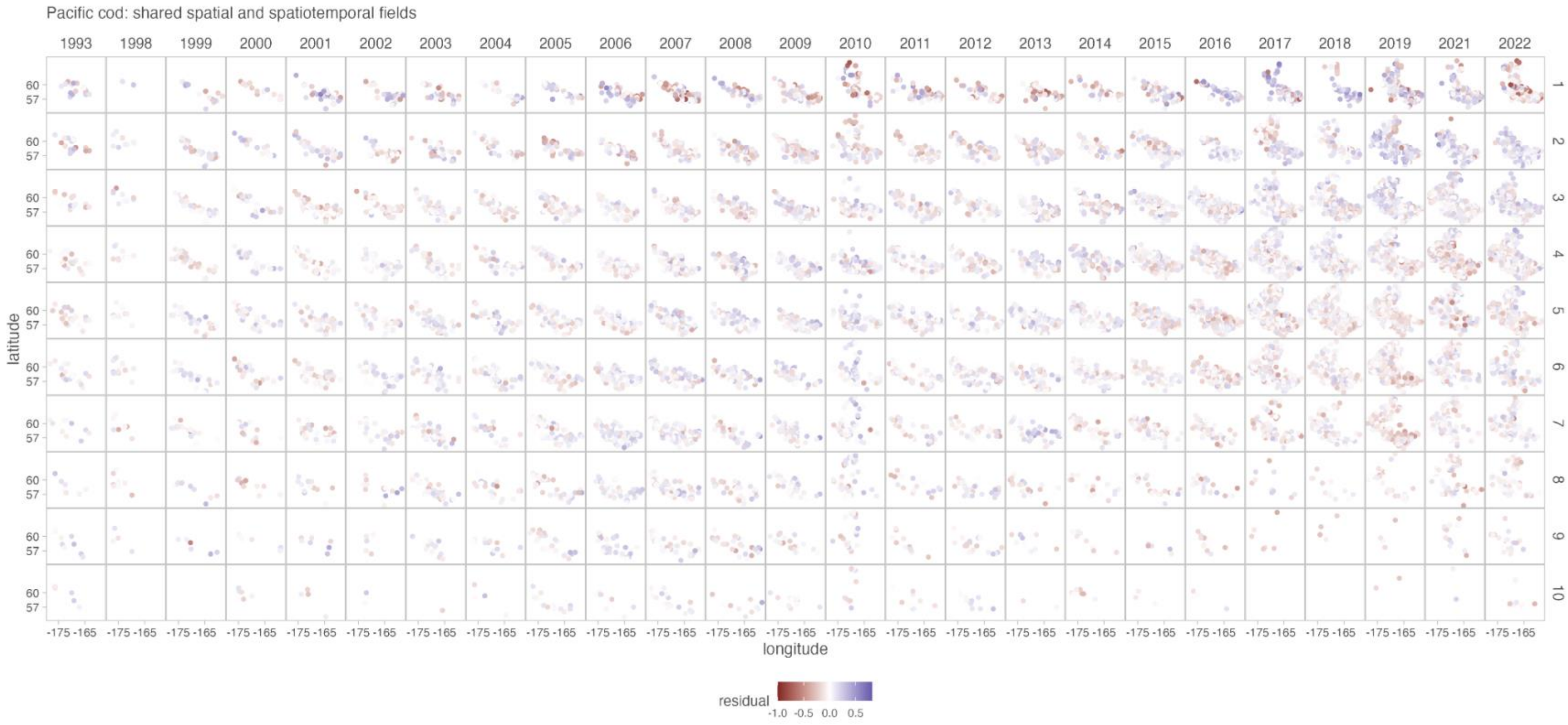

Pacific cod: shared spatial and spatiotemporal fields
1993
1998
1999
2000
2001
2002
2003
2004
2005
2006
2007
2008
2009
2010
2011
2012
2013
2014
2015
2016
2017
2018
2019
2021
2022
1
2
3
4
5
6
7
8
9
10
60
57
latitude
-175 -165
longitude
residual
-1.0 -0.5 0.0 0.5

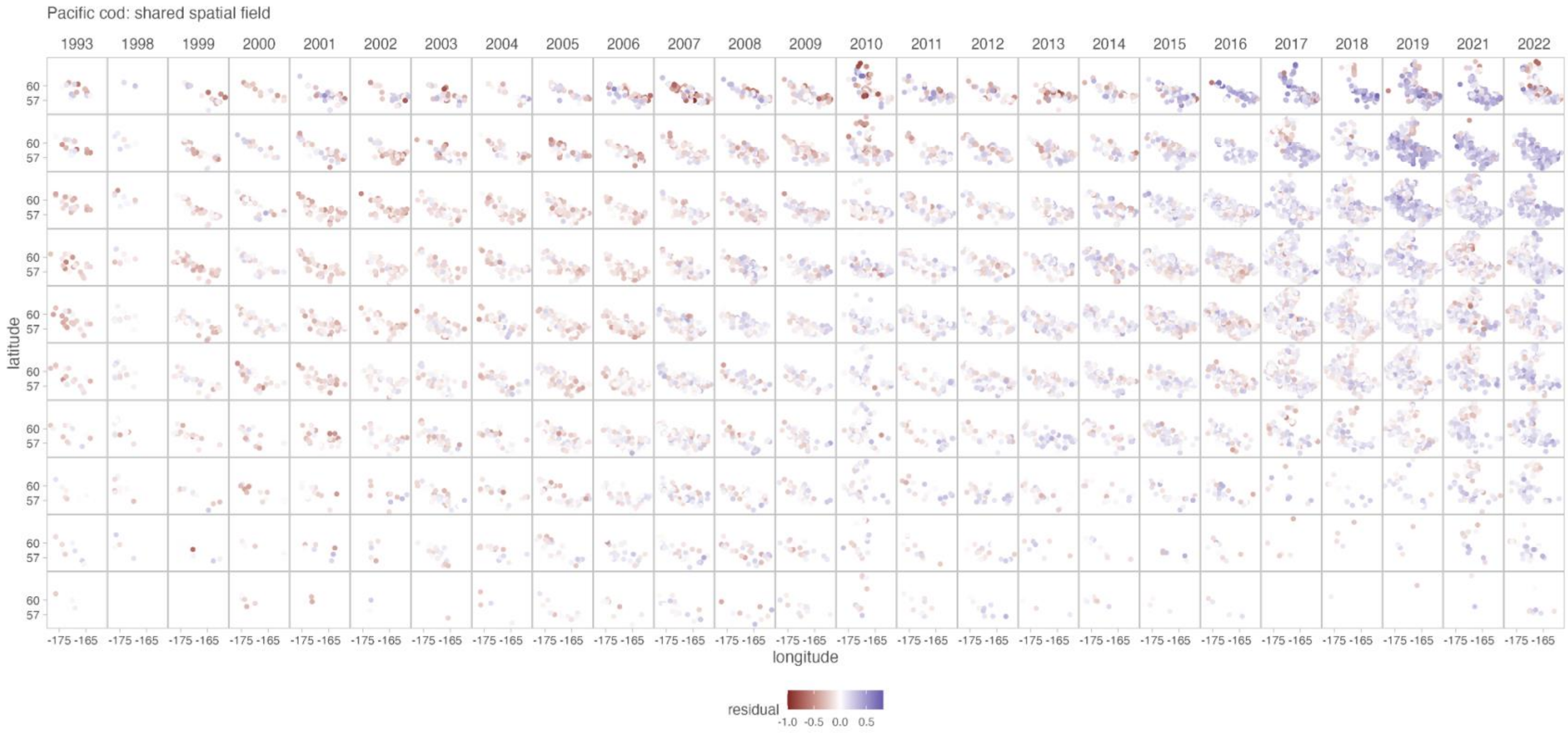

Figure S7. Comparison of residuals over space and time for Pacific cod among models with the same fixed effect structure but the three different random effect structures (see text): age-specific fields (first figure), shared spatial and spatiotemporal fields (second figure), and shared spatial fields (third figure). Residuals shown here are from the most supported model with the most supported random effect structure, i.e., the most supported model in Table S7, which has a second order polynomial relationship between weight and temperature with no interaction between age and temperature. Age class is indicated by row (i.e., 2 = Age 2).

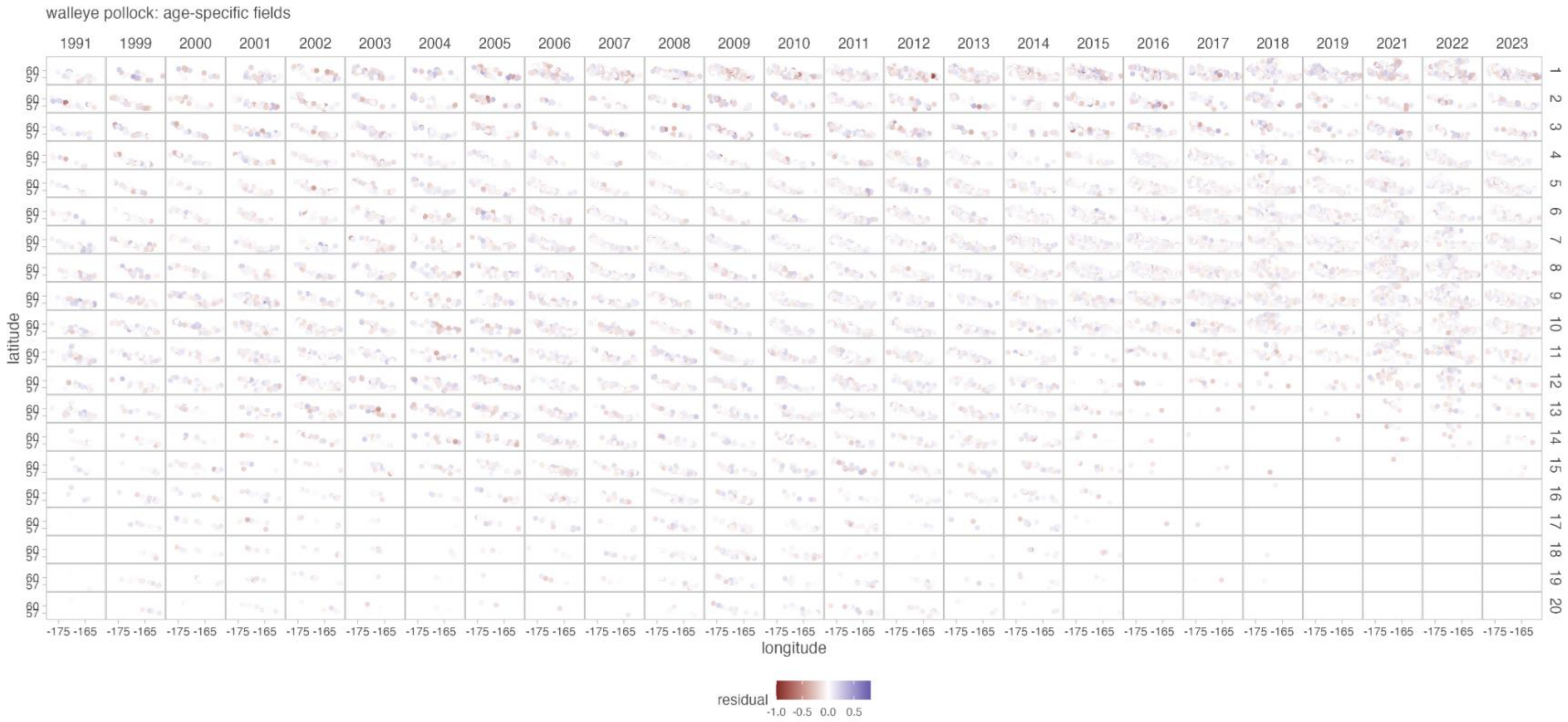

walleye pollock: age-specific fields
1991
1999
2000
2001
2002
2003
2004
2005
2006
2007
2008
2009
2010
2011
2012
2013
2014
2015
2016
2017
2018
2019
2021
2022
2023
latitude
60
57
-175 -165
longitude
residual
-1.0
-0.5
0.0
0.5

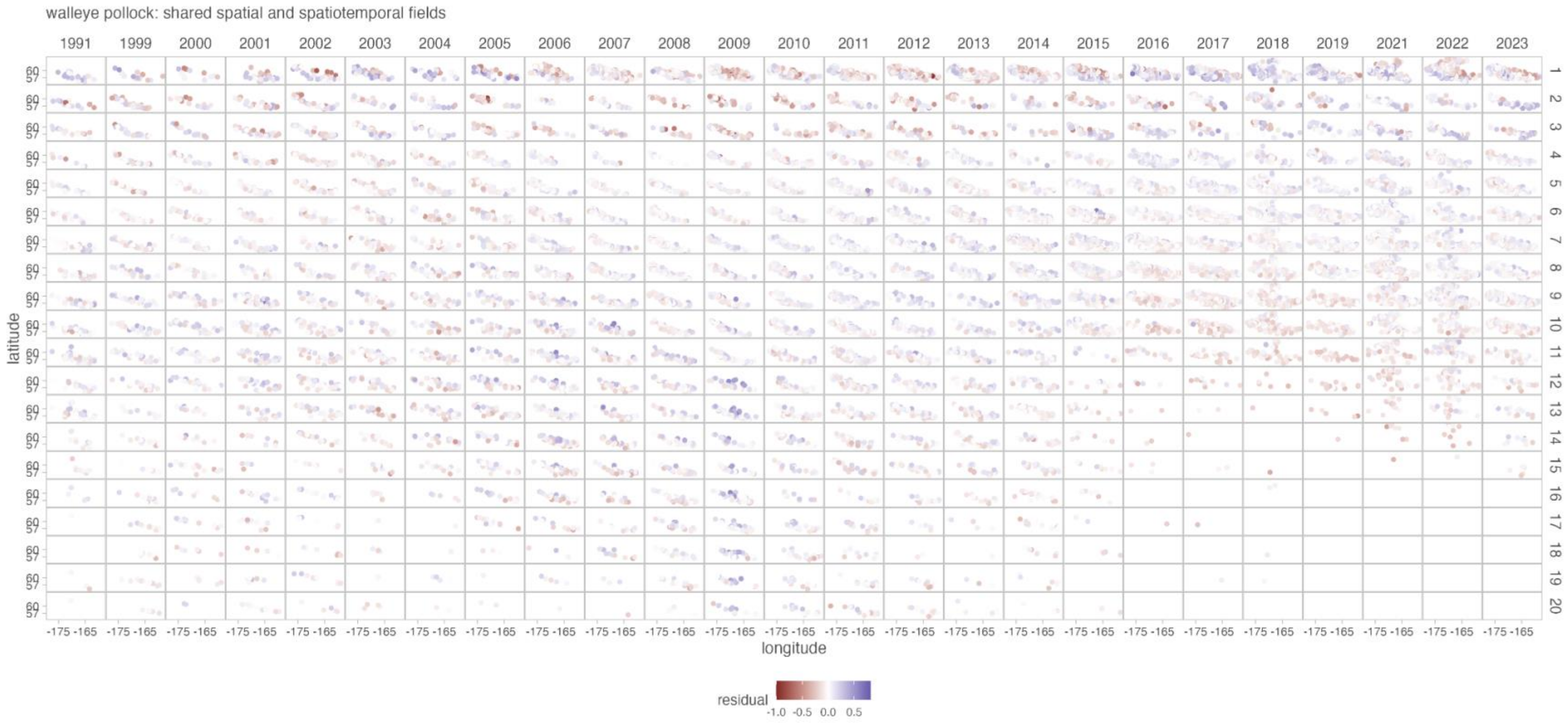

walleye pollock: shared spatial and spatiotemporal fields
1991
1999
2000
2001
2002
2003
2004
2005
2006
2007
2008
2009
2010
2011
2012
2013
2014
2015
2016
2017
2018
2019
2021
2022
2023
latitude
60
57
longitude
-175 -165
residual
-1.0
-0.5
0.0
0.5

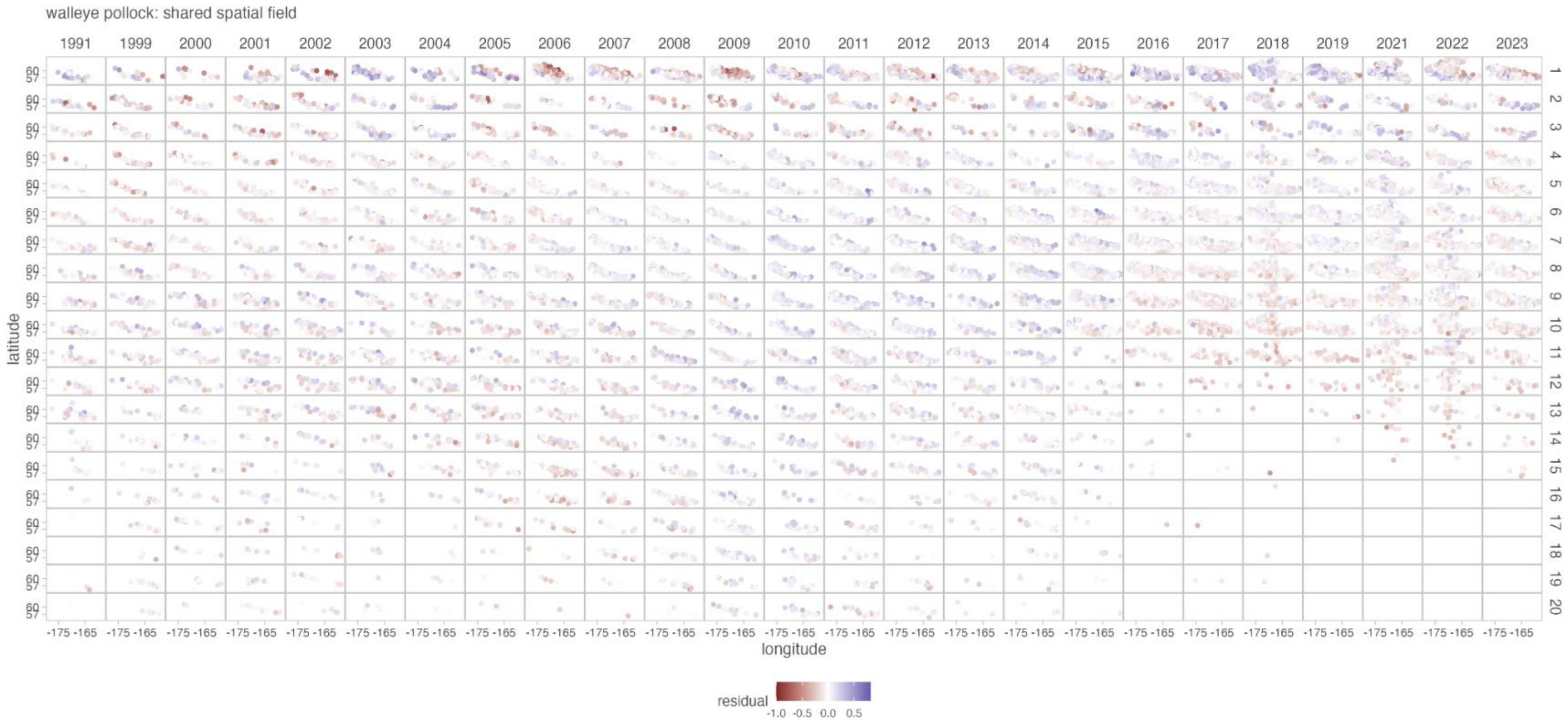

Figure S8. Comparison of residuals over space and time for walleye pollock among models with the same fixed effect structure but the three different random effect structures (see text): age-specific fields (first figure), shared spatial and spatiotemporal fields (second figure), and shared spatial fields (third figure). Residuals shown here are from the most supported model with the most supported random effect structure, i.e., the most supported model in Table S8, which has a second order polynomial relationship between weight and temperature with no interaction between age and temperature. Age class is indicated by row (i.e., 2 = Age 2).

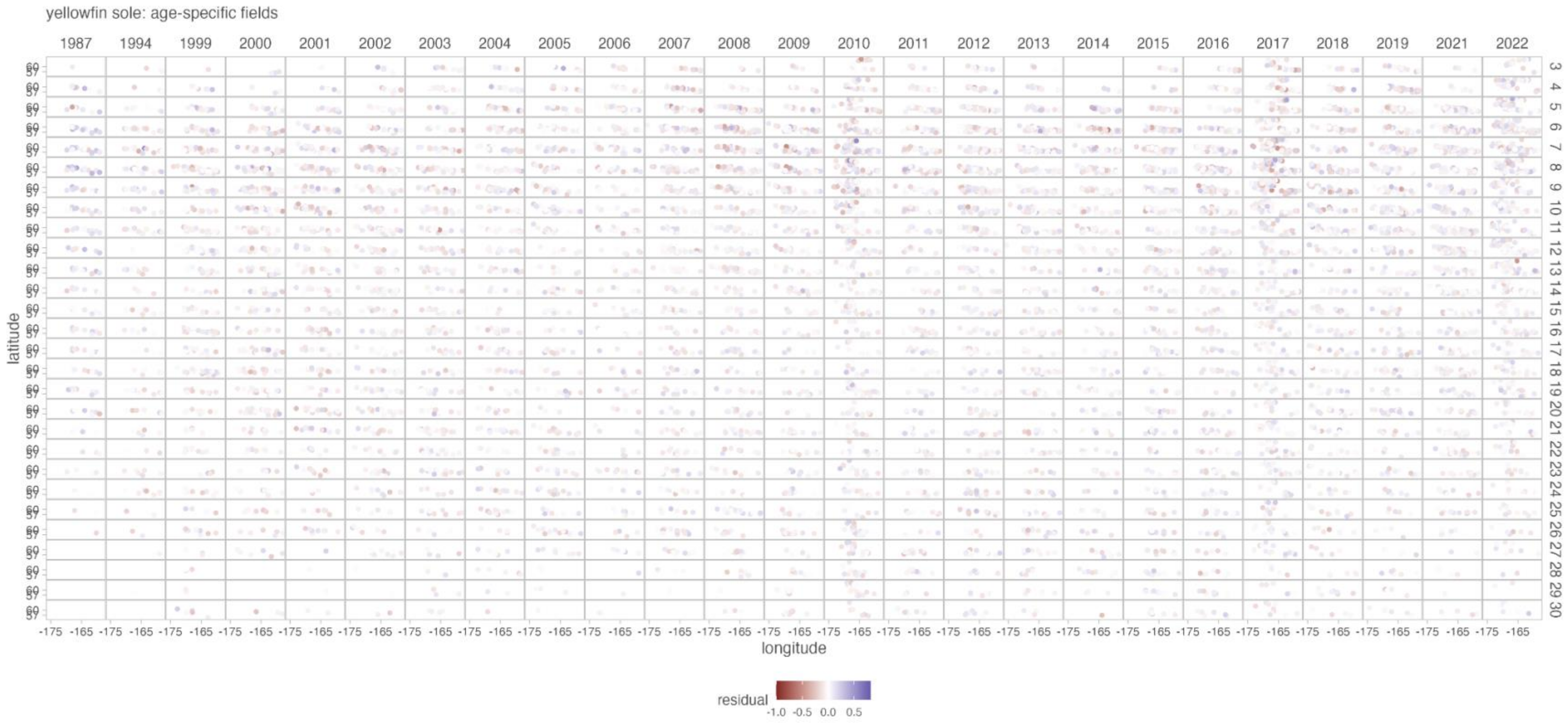

yellowfin sole: age-specific fields
1987
1994
1999
2000
2001
2002
2003
2004
2005
2006
2007
2008
2009
2010
2011
2012
2013
2014
2015
2016
2017
2018
2019
2021
2022
latitude
longitude
residual
-1.0 -0.5 0.0 0.5

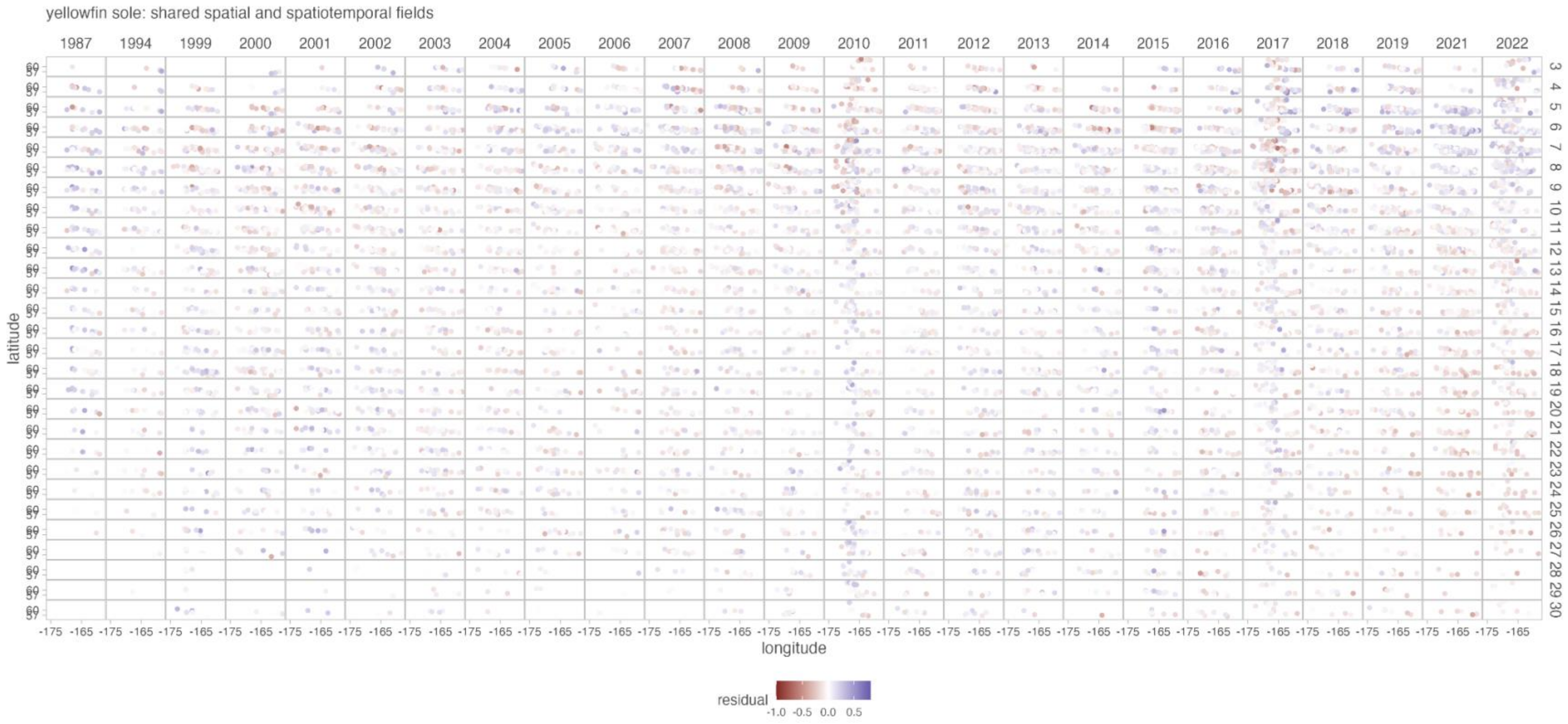
yellowfin sole: shared spatial and spatiotemporal fields
1987
1994
1999
2000
2001
2002
2003
2004
2005
2006
2007
2008
2009
2010
2011
2012
2013
2014
2015
2016
2017
2018
2019
2021
2022
latitude
longitude
residual
-1.0
-0.5
0.0
0.5

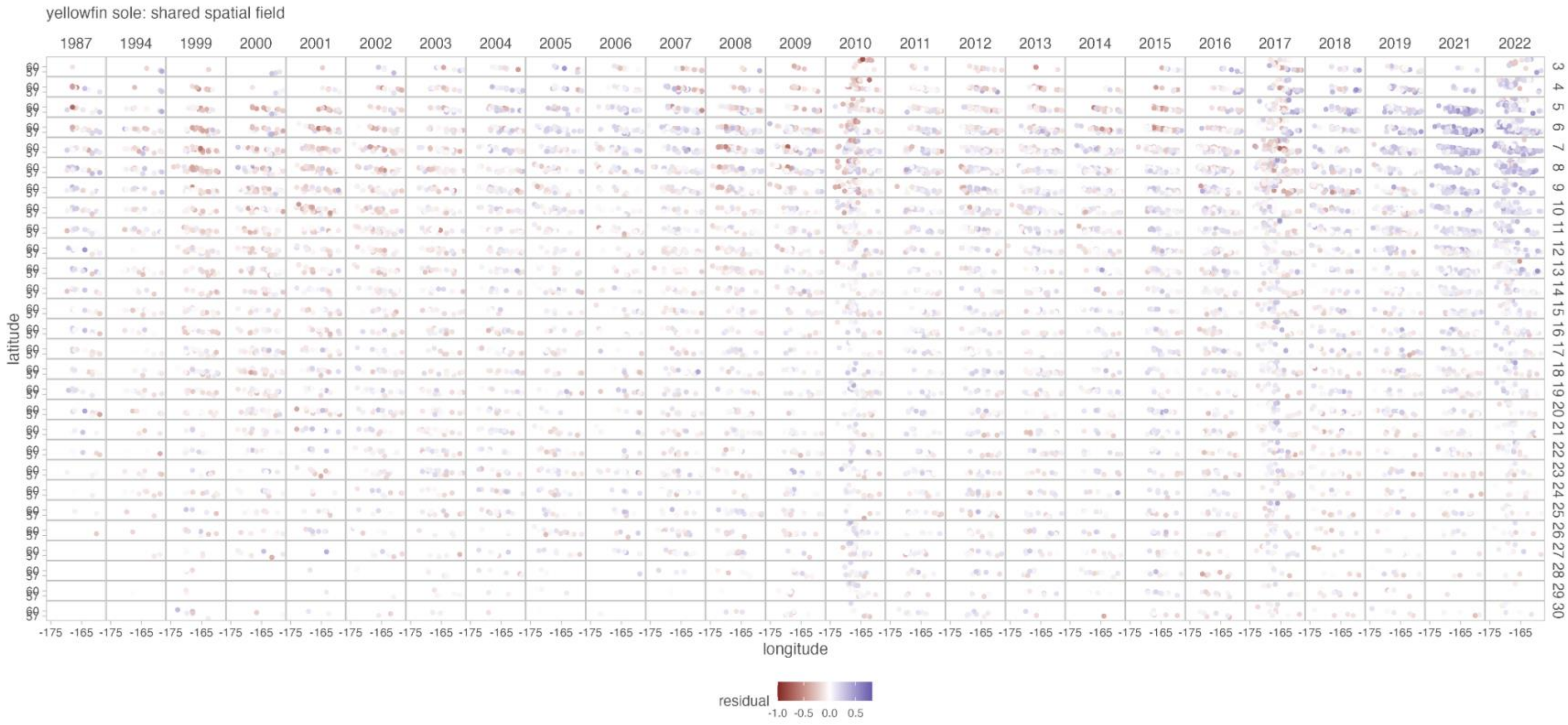

Figure S9. Comparison of residuals over space and time for yellowfin sole among models with the same fixed effect structure but the three different random effect structures (see text): age-specific fields (first figure), shared spatial and spatiotemporal fields (second figure), and shared spatial fields (third figure). Residuals shown here are from the most supported model with the most supported random effect structure, i.e., the most supported model in Table S9, which has a second order polynomial relationship between weight and temperature with no interaction between age and temperature. Age class is indicated by row (i.e., 2 = Age 2).

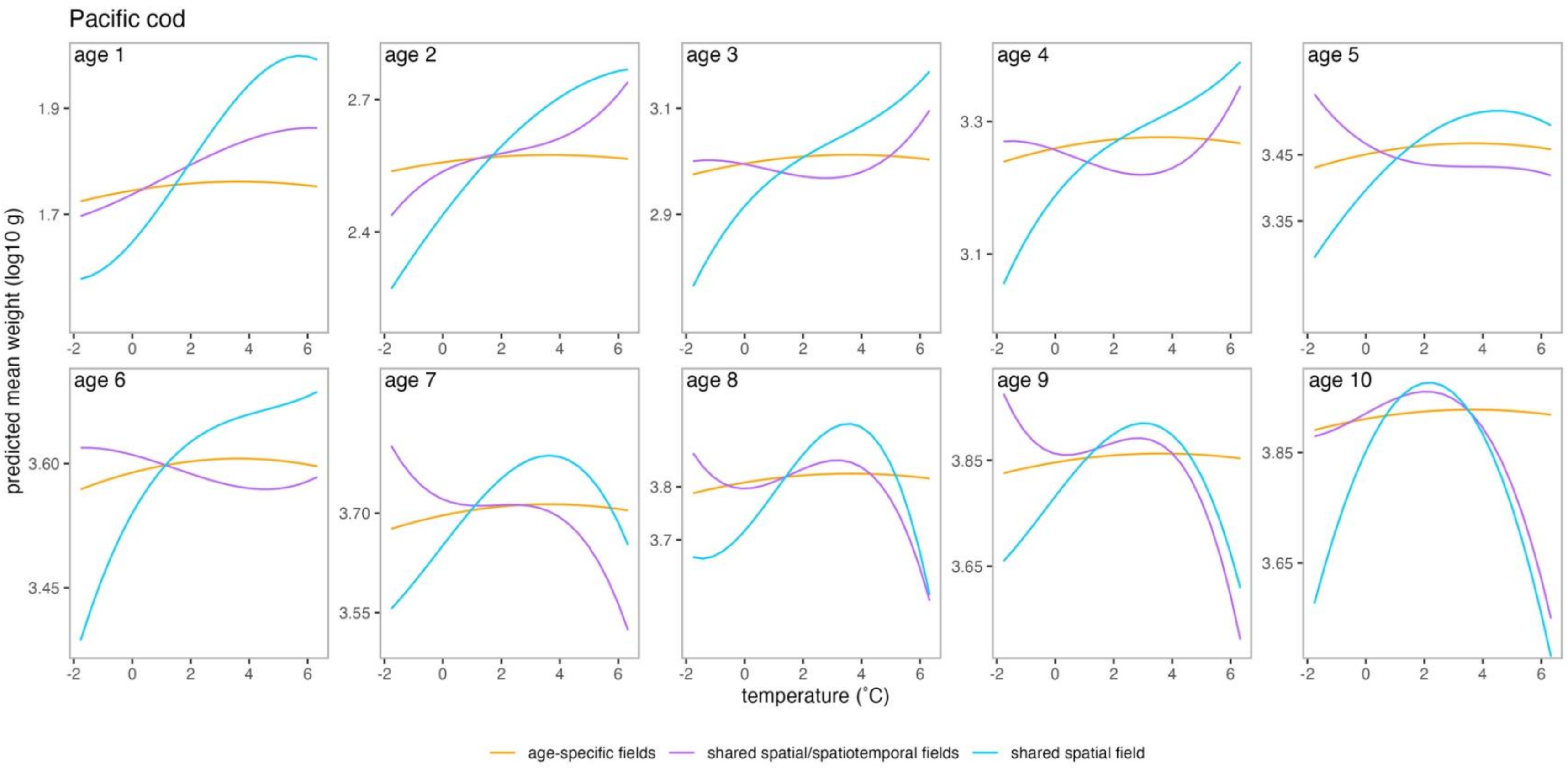

Figure S10. Predictions of Pacific cod weight from temperature from three spatiotemporal models: (1) age-specific spatial and spatiotemporal fields (orange lines), (2) shared spatial field and spatiotemporal fields across ages (purple lines), and (3) shared spatial field across ages with no spatiotemporal fields (blue lines) using data from years 2008 and forward (Figure 3 in main text has results with full dataset). Each panel corresponds to a separate age class (labeled in the upper left-hand side of the panel). Predictions in figure for the three models with different random effect structures (orange, purple and blue lines) were generated from the most supported model for each random effect structure (i.e., most supported models for each random effect structure in Table S3). Note the y-axis (the predicted mean weight [log10 g]) differs for each panel.
